# GalNAc-siRNA Mediated Knockdown of Ketohexokinase Versus Systemic, Small Molecule Inhibition of its Kinase Activity Exert Divergent Effects on Hepatic Metabolism in Mice on a HFD

**DOI:** 10.1101/2023.08.14.553218

**Authors:** Se-Hyung Park, Taghreed Fadhul, Lindsey R. Conroy, Harrison Clarke, Ramon C. Sun, Kristina Wallenius, Jeremie Boucher, Gavin O’Mahony, Alessandro Boianelli, Marie Persson, Genesee J. Martinez, Terry D. Hinds, Senad Divanovic, Samir Softic

## Abstract

Consumption of diets high in sugar and fat are well-established risk factors for the development of obesity and its metabolic complications, including non-alcoholic fatty liver disease. Metabolic dysfunction associated with sugar intake is dependent on fructose metabolism via ketohexokinase (KHK). Here, we compared the effects of systemic, small molecule inhibition of KHK enzymatic activity to hepatocyte-specific, GalNAc-siRNA mediated knockdown of KHK in mice on a HFD. Both modalities led to an improvement in liver steatosis, however, via substantially different mechanisms. KHK knockdown profoundly decreased lipogenesis, while the inhibitor increased the fatty acid oxidation pathway. Moreover, hepatocyte-specific KHK knockdown completely prevented hepatic fructose metabolism and improved glucose tolerance. Conversely, KHK inhibitor only partially reduced fructose metabolism, but it also decreased downstream triokinase. This led to the accumulation of fructose-1 phosphate, resulting in glycogen accumulation, hepatomegaly, and impaired glucose tolerance. In summary, KHK profoundly impacts hepatic metabolism, likely via both kinase-dependent and independent mechanisms.

**HIGHLIGHTS:**

- KHK knockdown or inhibition of its kinase activity differently target hepatic metabolism.
- KHK inhibitor increases F1P and glycogen accumulation as it also lowers triokinase.
- KHK knockdown completely prevents hepatic fructose metabolism and lipogenesis.
- E of wild type, but not mutant, kinase dead KHK-C increases glycogen accumulation.

## INTRODUCTION

Consumption of a calorie-dense high-fat diet (HFD) had long been considered the driver of the worldwide epidemic of obesity and its metabolic complications, including non-alcoholic fatty liver disease (NAFLD). A recent paradigm shift has brought more attention to dietary sugar as a major culprit fueling the obesity epidemic. Therefore, the World Health Organization (Nishida et al., 2004) and several leading medical societies, such as the American Heart Association (Johnson et al., 2009), Canadian Diabetes Association (Canadian Diabetes Association Clinical Practice Guidelines Expert et al., 2013) and three European medical groups (EASL, EASD, EASO) (European Association for the Study of the et al., 2016) recommend reducing sugar intake as a way to decrease obesity-associated complications.

The adverse effects of sugar have been mainly attributed to its fructose component. Several properties of dietary fructose make it uniquely suitable to promote the development of obesity-associated complications, such as NAFLD (Softic and Kahn, 2019). Fructose strongly enhances hepatic de novo lipogenesis (DNL) by acting as both a substrate for fatty acid synthesis (Zhao et al., 2020) and by upregulating SREBP1c and ChREBP transcription factors in the liver to elevate the expression of DNL enzymes (Softic et al., 2016b; Softic et al., 2017). On the other hand, fructose restriction improves hepatic steatosis (Cohen et al., 2021) and lowers body mass index (Radulescu et al., 2022). Furthermore, fructose decreases fatty acid oxidation (FAO) (Inci et al., 2022) either indirectly through DNL intermediate malonyl-CoA, which inhibits carnitine palmitoyltransferase 1a (CPT1α), the rate limiting enzyme of FAO, or directly through suppression of CPT1α by ketohexokinase (KHK), the first enzyme of fructolysis (Softic et al., 2019). Additionally, fructose has been proposed to induce hepatic insulin resistance (Softic et al., 2020), elevate uric acid production (Lanaspa et al., 2012), promote endoplasmic reticulum stress (Lee et al., 2008), and propagate mitochondrial dysfunction (Chiang Morales et al., 2022), all pathways that contribute to metabolic impairment. Indeed, fructose consumption has been found to be 2-3 fold higher in adults with biopsy-confirmed NAFLD, as compared to BMI-matched controls (Ouyang et al., 2008). Moreover, adults with increased fructose consumption have a higher incidence of liver fibrosis (Abdelmalek et al., 2010).

The effects of dietary fructose are dependent on its catabolism via KHK. Thus, mice with whole body KHK knockout on an obesogenic diet do not develop NAFLD (Ishimoto et al., 2012; Ishimoto et al., 2013). We and others have shown that deletion of KHK, specifically in the liver, is sufficient to reverse NAFLD (Andres-Hernando et al., 2020; Softic et al., 2017). Based on a strong premise that fructose contributes to metabolic dysfunction and that knockout of KHK prevents it has led to the development of small molecule inhibitors of KHK enzymatic activity. The inhibitors target the ATP binding domain of KHK, which prevents fructose phosphorylation to fructose-1 phosphate (F1P) (Maryanoff et al., 2012). Phosphorylation of fructose, similar to phosphorylation of glucose, is required for its downstream metabolism. An early KHK inhibitor (Maryanoff et al., 2011) was efficacious in preclinical studies, but this compound was never tested in clinical trials. Many enzymes contain an ATP binding domain, so the initial inhibitors were not selective enough for KHK activity. Within the last decade, a new class of compounds, allegedly 600 times more specific for KHK activity, have been developed (Futatsugi et al., 2020; Huard et al., 2017). These inhibitors were likewise found to effectively reduce steatosis in rats (Gutierrez et al., 2021) but also in human liver tissue (Shepherd et al., 2021). Moreover, recent clinical trials reported that the new KHK inhibitors significantly reduced whole liver fat in patients with NAFLD (Kazierad et al., 2021; Saxena et al., 2022). In spite of the positive data, the leading developer unexpectedly announced that they are stopping further advancement of KHK inhibitors (Taylor, 2021).

In this study, we compared the effects of systemic, small molecule KHK inhibitor (PF-06835919) versus hepatocyte-specific KHK knockdown (KD) using GalNAc conjugated siRNA targeting mouse total KHK. We found that both KHK KD and inhibition of its kinase activity effectively reduced fat accumulation in the liver. However, this outcome is achieved through distinctly different effects on hepatic metabolism. KHK siRNA reduced the hepatic DNL pathway resulting in lower steatosis and improved glucose tolerance. On the other hand, the KHK inhibitor decreased steatosis by upregulating the FAO pathway. Additionally, the KHK inhibitor only partially reduced liver fructose metabolism, but unexpectedly it also lowered downstream triokinase (TKFC), resulting in increased F1P accumulation and glycogen deposition in the liver. Our laboratory was the first to suggest that KHK may have kinase independent functions (Helsley et al., 2023; Park et al., 2023). Here we show that inhibiting KHK kinase activity leads to different metabolic outcomes compared to a complete deletion of the KHK protein.

## RESULTS

### Inhibition of ketohexokinase activity, but not its knockdown, leads to hepatic glycogen accumulation

To compare the effects of liver-specific KHK knockdown using siRNA versus systemic small molecule inhibitor of KHK kinase activity we placed male, C57Bl6/J, mice on a low-fat diet (LFD) or a high-fat diet for six weeks. Thereafter, the mice on high-fat diet were subdivided into three groups; control group (HFD), KHK siRNA injected group (HFD+siRNA) and KHK inhibitor gavaged group (HFD+Inhib). All mice were injected with luciferase control siRNA (10 mg/kg) or siRNA targeting total KHK (20mg/kg) every two weeks. Also, all mice were gavaged methylcellulose control or (15mg/kg) KHK inhibitor (PF-06835919) in methylcellulose, twice daily for four weeks (**Fig S1A**). The dose of the inhibitor was set to achieve an exposure in mice that has been previously shown to achieve high KHK inhibition in rats and humans (Gutierrez et al., 2021). The mice were treated with the drugs for four weeks and sacrificed. The mice in the HFD group weighed, significantly more than the mice on a LFD (37.8±1.1g vs. 27.5±0.9g) (**Figs 1A, S1B**). KD of KHK (36.3±0.9g) or inhibition of its kinase activity (35.5±0.8g) did not significantly reduce body weight. In agreement with weight gain, caloric intake was increased in all mice on a HFD compared to the LFD control, but there was no difference among the groups on HFD (**Fig S1C**). Similarly, water intake was not different among the groups on a HFD (**Fig S1D**). Body composition assessed by echoMRI revealed lower lean mass percent in HFD (66.8±3.3%), compared to the LFD group (85.4±1.2%) (**Fig 1B**). KD of KHK resulted in no change (65.1±1.4%), while inhibition of KHK increased percent lean mass (72.9±1.8%). Conversely, percent body fat was higher in mice on a HFD (30.5%±3.2), compared to the LFD group (11.3±1.3%). The HFD+siRNA group (32.6±1.4%) was not different, while percentage of body fat decreased in the HFD+Inhib group (24.2±2.0%), which could be accounted by lower perigonadal adipose tissue weight (**Fig 1C**), while the increased lean mass in the HFD+Inhib group likely reflects increased liver weight in these mice (**Fig 1D**). In addition to liver, kidney and intestines also effectively metabolize fructose via KHK (Helsley et al., 2020). Kidney weight was not altered (**Fig S1E**) and histology of the intestines was unchanged among the groups (**Fig S1F**).

**Figure 1.**
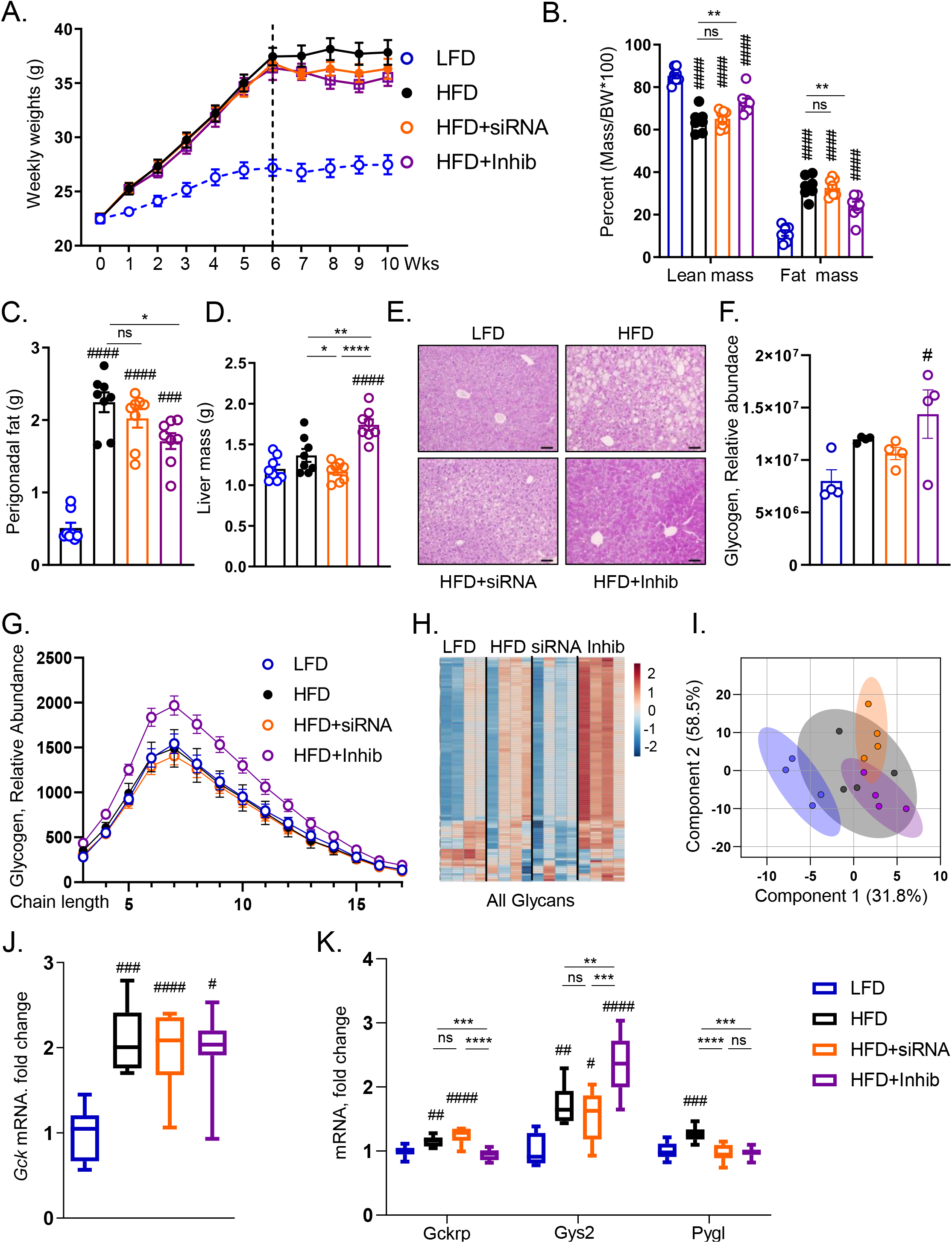
Inhibition of ketohexokinase activity, but not its knockdown, leads to hepatic glycogen accumulation. (A) Weight gain of mice on low-fat diet (LFD), high-fat diet (HFD), HFD treated with siRNA, and HFD treated with inhibitor for the last four weeks of this ten weeks experiment. (B) Lean mass and fat mass normalized by body weight as assessed my EchoMRI. Perigonadal adipose tissue (C) and liver (D) weights at the time of sacrifice. n=7-8 mice per group. (E) Representative periodic acid-schiff (PAS) stained images of liver histology. Bar=50µm. (F) Mass spectrometry (MS) analysis for glycogen in the liver. (G) Glycogen chain length as determined by MS. (H) Heatmap of all glycans and (I) principal component analysis of all glycans in LFD, HFD, HFD+siRNA, and HFD+Inhib groups. n=4 mice per group. mRNA expression of Gck (J) and the genes involved in glycogen synthesis and degradation. n=6 mice per group. (K). Statistical analysis was performed using one way ANOVA compared to LFD group (#p < 0.05; ## p < 0.01; ### p < 0.001; #### p < 0.0001) with post hoc t tests between the individual groups (*p < 0.05; **p < 0.01; ***p < 0.001; ****p < 0.0001).

A surprising increase in liver weight in the HFD+Inhib group could not be explained by liver steatosis (**Fig S1G**), as liver triglycerides decreased in both HFD+siRNA and HFD+Inhib groups, compared to the HFD (**Fig S1H**). On the other hand, the periodic acid-Schiff (PAS) stain suggested that glycogen amount was higher in the HFD+Inhib group (**Fig 1E**). Indeed, mass spectrometry quantification revealed elevated glycogen levels only in the HFD+Inhib group, compared to the LFD group (**Fig 1F**). Glycogen accumulation is controlled by glucagon and indeed we found reduced glucagon concentration in HFD+Inhib group (39±4pg/ml), compared to both HFD (63±7pg/ml) and HFD+siRNA groups (62±6 pg/ml) (**Fig S1I**). Next, we quantified glycogen chain length and found that only HFD+Inhib group had an increased abundance of glycogen composed of short glucose polymers (**Fig 1G**). Short chain glycogen more readily releases glucose, so next we quantified the levels of all N-linked glycans. Glycans are homo- or heteropolymers of monosaccharide residues attached to a protein (Conroy et al., 2021). A heatmap representation of all glycans documented that glycans were higher in the HFD, compared to the LFD group. Glycans were decreased in the HFD+siRNA, compared to the HFD group, but were the highest in the HFD+Inhib group (**Fig 1H**), indicative of greater monosaccharide availability. The principal component analysis of all glycans confirmed differential clustering in HFD versus LFD groups (**Fig 1I**). On a HFD, KD of KHK clustered glycans in a new direction, while inhibition of KHK activity affected clustering in a unique way. Together these data indicate that mice in the HFD+Inhib group had the highest accumulation of glycogen in the liver and the greatest monosaccharide availability.

To interrogate the mechanism behind increased glycogen accumulation we quantified the expression of glucokinase (Gck), which was increased two-fold in all mice on high-fat diet, but was not altered by KHK KD or inhibition (**Fig 1J**). In its inactive state, Gck is sequestered in the nucleus by glucokinase regulatory protein (Gckrp) (Sternisha and Miller, 2019). Upon disassociation from Gckrp, Gck migrates into the cytoplasm where it converts glucose to glucose-6 phosphate to be used for glycogen synthesis. The mice in the HFD and HFD+siRNA groups had 1.3 to 1.4-fold higher Gckrp expression compared to the LFD group (**Fig 1K**). The HFD+Inhib group did not have elevated Gckrp mRNA, permitting Gck to be more active. Gck activity stimulates glycogen synthase (Gys2), which was 1.5 to 1.7-fold higher in HFD and HFD+siRNA groups and 2.4-fold higher in the HFD+inhib group, compared to the LFD control. Conversely, glycogen phosphorylase, which liberates glucose from glycogen, was only elevated in mice on a HFD, compared to LFD, but was not affected in the HFD+siRNA or HFD+Inhib groups. In summary, both knockdown and inhibition of KHK improves hepatic steatosis, but KHK inhibition increased liver weight, in part, due to higher glycogen accumulation.

### Knockdown of ketohexokinase, but not inhibition of its kinase activity, improves glucose tolerance and increases the glycolysis pathway

As expected, the mice on a HFD had impaired glucose tolerance compared to the LFD-fed mice (**Figs 2A, 2B**). KD of KHK improved glucose tolerance, but inhibition of KHK kinase activity did not. Similarly, fasted blood glucose was elevated in mice on a HFD (171±9mg/dL) compared to the LFD group (132±3mg/dL) (**Fig 2C**). Glucose improved following KHK KD (138±5mg/dL), but not with inhibition of its kinase activity (157±6mg/dL). Fasted insulin was higher in the HFD (2.0±0.2ng/ml) compared to the LFD group (0.3±0.1ng/ml), and was reduced in both KHK KD (0.9±0.1ng/ml) and the inhibitor groups (0.8±0.1ng/ml) (**Fig 2D**). Similarly, HOMA-IR, a measure of whole-body insulin resistance, was elevated in mice on a HFD and was decreased with both KHK KD and inhibition of its kinase activity (**Fig 2E**). Insulin signaling in the liver was assessed by injecting insulin or saline via the inferior vena cava 10 minutes before sacrifice. Insulin-stimulated Akt phosphorylation was decreased in mice on HFD compared to the LFD group (**Fig 2F**). KD of KHK and inhibition of its activity both improved Akt phosphorylation, in agreement with reduced hepatic steatosis in these mice. Erk phosphorylation was also decreased in the HFD compared to the LFD group, but it did not improve with KD or inhibition of KHK. Thus, improved glucose tolerance following KHK KD is not due to improved hepatic insulin signaling, which was equally better with both KD of KHK or inhibition of KHK activity.

**Figure 2.**
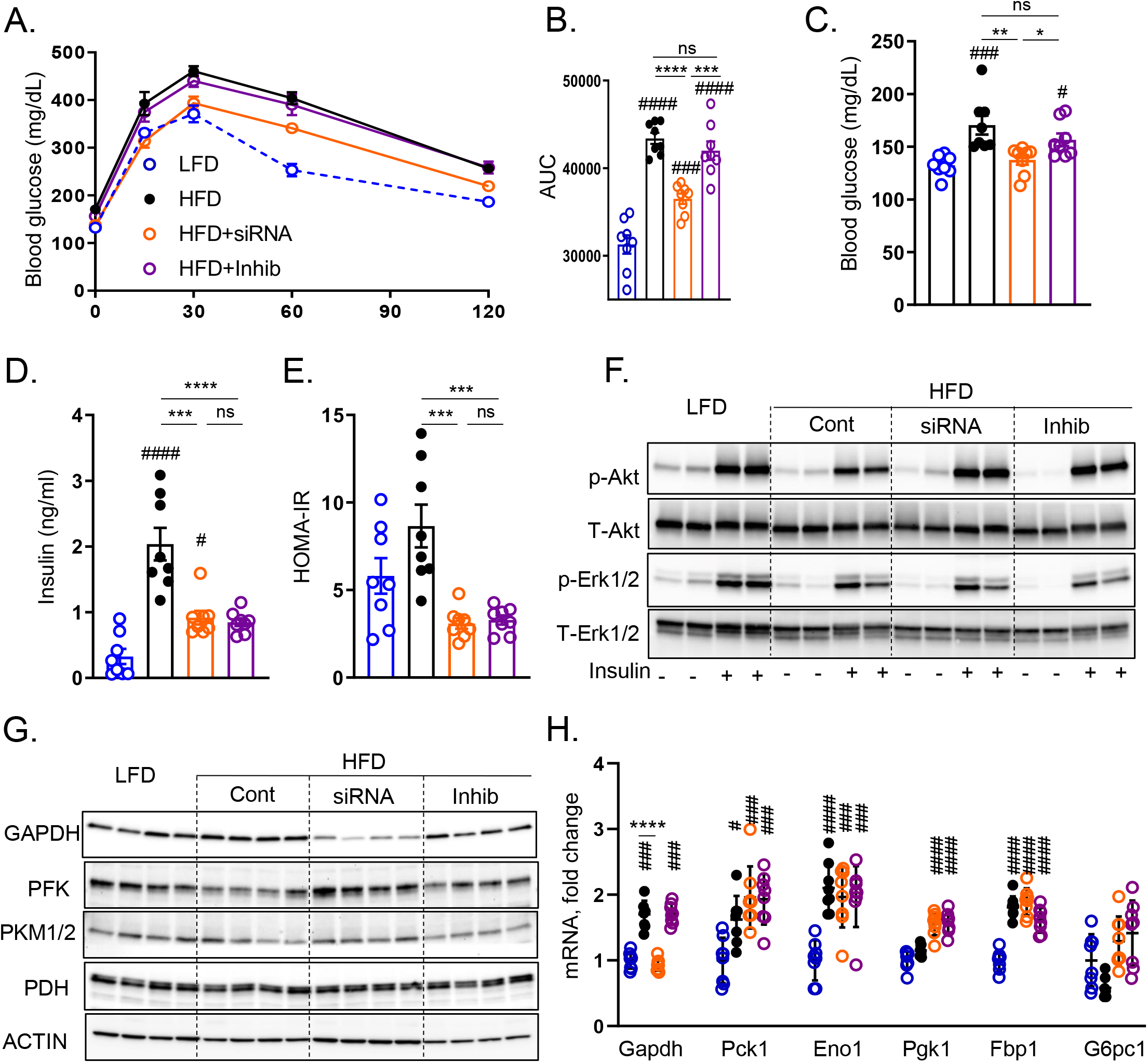
Knockdown of ketohexokinase, but not inhibition of its activity, improves glucose tolerance and increases the glycolysis pathway. (A) Glucose tolerance test (GTT) measured after 8 weeks on the diets and two weeks after initiation of the treatments. (B) Area under the curve calculated from GTT. n=7-8 mice per group. Fasted serum glucose (C), insulin (D), and calculated HOMA-IR (E) at 10 weeks on the diet (F) Western blot analysis of insulin signaling in the liver. n=4 mice per group. Western blot analysis (G) and qPCR quantification of mRNA (H) of genes mediating the gluconeogenesis pathway. n=6 mice per group for gene expression. Actin was used as a loading control. Statistical analysis was performed using one way ANOVA compared to LFD group (### p < 0.001; #### p < 0.0001) with post hoc t tests between the individual groups (*p < 0.05; **p < 0.01; ***p < 0.001; ****p < 0.0001).

Next, we assessed the glucose utilization pathway by Western blot. Glyceraldehyde-3-phosphate dehydrogenase (GAPDH) controls the flow of fructose carbons onto the glycolysis pathway. GAPDH was slightly increased in the HFD-compared to the LFD-fed mice (**Figs 2G, S2**). KHK KD profoundly decreased GAPDH, in agreement with the abrogation of fructose metabolism. Inhibition of KHK activity did not result in lower GAPDH. The rate limiting enzyme of glycolysis, phosphofructokinase (PFK), was decreased in the HFD group, compared to the LFD group (**Figs 2G, S2**). The mice in HFD+siKHK group had elevated PFK protein compared to the HFD group indicative of increased glycolysis. On the other hand, the mice in HFD+Inhib group did not have elevated PFK. The last enzyme of glycolysis, pyruvate kinase (PK) is allosterically activated by fructose-1,6-bisphosphate, the product of PFK activity. Therefore, PK was elevated in HFD+siRNA group, compared to the HFD+inhib group. Pyruvate dehydrogenase (PDH) converts pyruvate into acetyl-CoA to be used in the citric acid cycle. PDH was unchanged among the groups. Together, these data demonstrate that glycolysis pathway is elevated in HFD+siRNA group, but not in HFD+Inhib group.

We also assessed the gluconeogenesis pathway by qPCR. Gapdh is an enzyme mediating the fructolysis and gluconeogenesis pathways, and its expression mimicked GAPDH protein, which likely reflects its role in the fructolysis pathway (**Fig 2H**). All other enzymes involved in gluconeogenesis, phosphoenolpyruvate carboxykinase 1 (Pck1), enolase (Eno1), phosphoglycerate kinase 1 (Pgk1), fructose-bisphosphatase 1 (Fbp1) and glucose-6-phosphatase (G6pc1) were not different between HFD+siKHK and HFD+Inhib groups. In summary, improved glucose tolerance in the HFD+siRNA, compared to the HFD+Inhib group is mediated via increased glycolysis pathway, but is not due to enhanced hepatic insulin signaling, which was improved in both groups.

### Small molecule inhibitor decreases ketohexokinase enzymatic activity in liver and kidney, but not in intestine

Due to a strong phenotypic difference between KHK KD and inhibition of its kinase activity we assessed how well these two modalities prevent hepatic fructose metabolism. Compared to the HFD-fed mice (549±17nM), serum fructose was elevated in HFD+Inhib treated mice (672±48nM) (**Fig 3A**), in line with systemic inhibition of fructose metabolism. Liver-specific KHK siRNA treated mice (580±31nM) did not have increased serum fructose, as fructose can also be metabolized in other tissue. As expected, liver fructose concentration was higher in both HFD+siRNA (30.0±2.8nmol/g) and HFD+Inhib (27.1±3.8nmol/g) groups, compared to the HFD group (16.1±1.6nmol/g) (**Fig 3B**). The product of KHK activity, F1P was significantly reduced in the HFD+siRNA group (6.2±1.3nmol/g), compared to the HFD group (12.7±1.8nmol/g) (**Fig 3C**). Surprisingly, the HFD+Inhib group (29.0±10.4nmol/g) had much higher hepatic F1P concentration, 2-5 hours after the inhibitor gavage. F1P is known to induce liver injury, so we quantified serum ALT. HFD group had higher ALT than LFD group, but ALT decreased in both HFD+siRNA and HFD+Inhib groups (**Fig S3A**). The expression of proinflammatory genes was also increased in the HFD, compared to the LFD group and was largely not affected by the two interventions, except for MCP1 which decreased in both HFD+siRNA and HFD+Inhib groups (**Fig S3B**).

**Figure 3.**
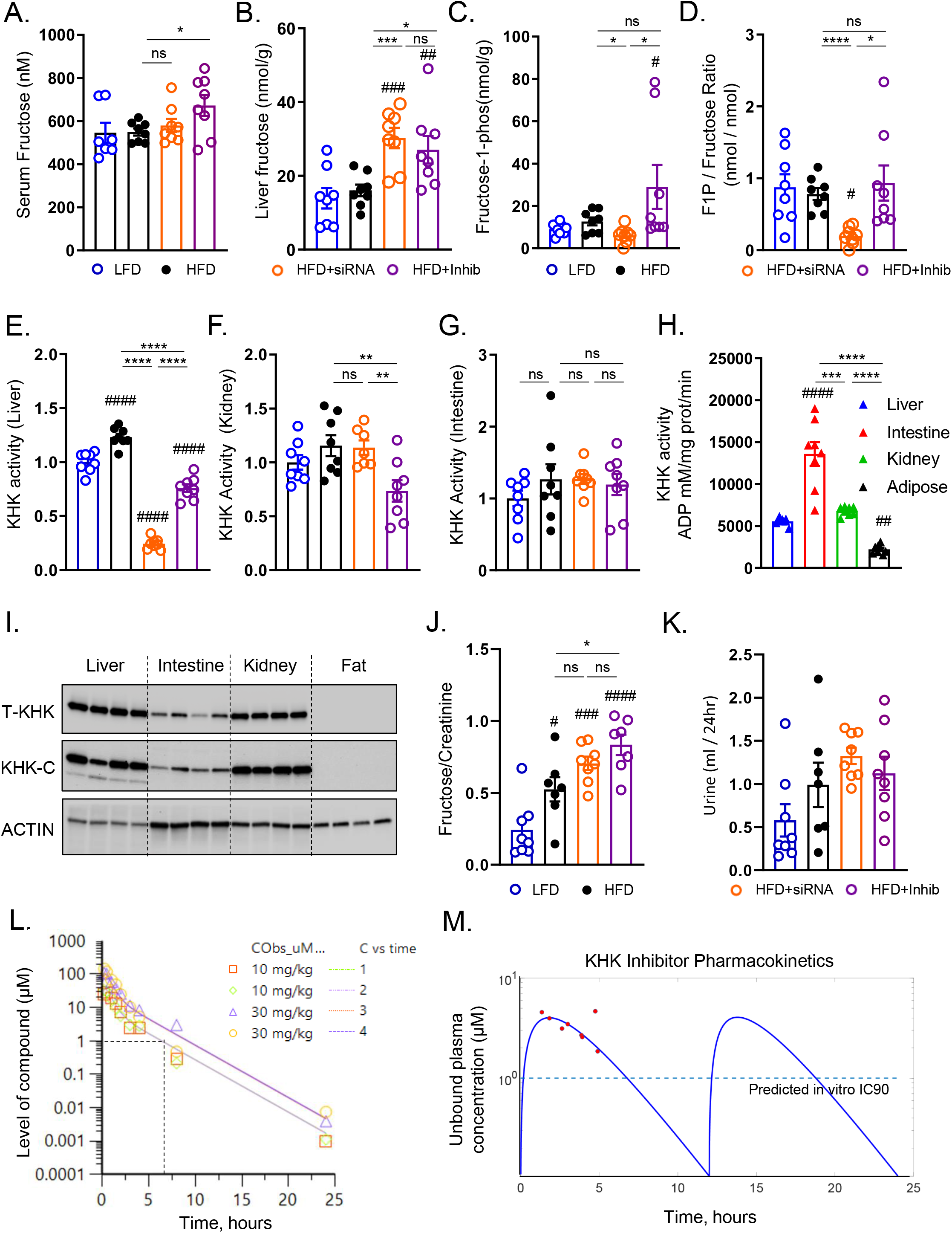
Small molecule inhibitor decreases ketohexokinase enzymatic activity in liver and kidney, but not in intestine. (A) Serum fructose level from the mice at sacrifice. The levels of (B) fructose (C), fructose 1-phosphate (F1P) and the ratio of (D) F1P/fructose in the liver. KHK activity in liver(G), kidney(H), and intestine(I) fed on LFD, HFD, HFD with siRNA, and HFD with inhibitor for 10 weeks. n=7-8 mice per group. Quantification of KHK activity in (E) liver, (F) kidney, and (G) intestine. Liver KHK activity is set to one in figures E-F. (H) Absolute KHK activity (K) in liver, intestine, kidney and perigonadal adipose tissue 2-5h after last inhibitor dosing. (I) Western blot of total KHK and KHK-C in liver, intestine, kidney and perigonadal adipose tissue. n=4 mice per group. Vinculin was used as a loading control. (J) Urinary fructose level corrected by urine creatinine and (K) urine volume collected for 24h. (L) *In vivo* monitoring of the inhibitor concentration over 24h period following single gavage with 10 mg/kg or 30 mg/ml of the inhibitor. n=2 mice in 10 and n=2 mice in 30 mg/kg dose. (M) Unbound Plasma concentration of the inhibitor (red dots) quantified by mass spectrometry in chow fed mice, 2-5h after last dose of the inhibitor. Pharmacokinetic model fitting (blue line) based on inhibitor concentration (red dots). Dashed line represents target concentration. Statistical analysis was performed using one way ANOVA compared to LFD group (#p < 0.05; ### p < 0.001; #### p < 0.0001) with post hoc t tests between the individual groups (*p < 0.05; **p < 0.01; ***p < 0.001; ****p < 0.0001).

Next, we calculated F1P to fructose ratio to approximate KHK activity. The ratio was reduced only in the KHK+siRNA group (**Fig 3D**), but it was equivalent in the HFD and HFD+Inhib groups, indicating that these mice can still metabolize fructose in the liver. KHK activity was also directly measured in liver homogenates utilizing our recently developed protocol (Park et al., 2021). The mice on a HFD had 23% higher KHK activity in the liver, compared to the mice on a LFD (**Fig 3E**). Compared to the HFD group, KD of KHK decreased KHK activity by 80%, likely reaching background activity levels of the assay. However, the inhibitor lowered KHK activity only by 38%. These data are in agreement with F1P to fructose ratio confirming that the inhibitor group can still metabolize fructose in the liver. In the kidney, only the inhibitor, but not siRNA treatment decreased KHK activity by 36%, in line with the systemic effects of the inhibitor (**Fig 3F**). However, in the intestine, both KHK+siRNA and KHK+Inhib groups did not induce lower KHK activity (**Fig 3G**). We compared KHK activity in the liver, intestine and kidney, to adipose tissue, which does not metabolize fructose and serves as a baseline. In LFD-fed mice, KHK activity was 2.4-fold higher in the intestine, as compared to both liver and kidney, while adipose tissue had background levels of KHK activity (**Fig 3H**). Next, we measured total KHK and KHK-C isoform in these tissues and found that the intestine has 3-times lower KHK-C protein compared to the liver and kidney, whereas adipose tissue does not express KHK (**Figs 3I, S3C**). These data are in agreement with a recent study documenting that the intestine metabolizes fructose more robustly than the liver, but it can only metabolize a small amount of fructose (Jang et al., 2018). Urine fructose normalized to creatinine was two-fold higher in HFD compared to LFD group (**Fig 3J**). The HFD+siRNA group did not show a further elevation in urinary fructose; however, the HFD+Inhib group had a stepwise higher urinary fructose. Urine creatinine was not different among the groups (**Fig S3D**). The volume of urine collected over 24 hours was higher in mice on HFD than on LFD, but was not affected by KHK KD or inhibition (**Fig 3K**). 24h urine fructose excretion was higher in mice on a HFD and it further increased by both KHK KD and inhibition of its activity (**Fig S3E**). In summary, KD of KHK resulted in an almost complete loss of fructose metabolism in the liver. On the other hand, systemic inhibition of KHK activity partially reduced fructose metabolism in liver and kidney, but not in intestine.

Due to differences in KHK activity and F1P/Fructose ratio, we tested if the dose of the inhibitor used was adequate to inhibit KHK activity *in vivo*. In a pilot study, the mice were gavaged 10 mg/kg, 30 mg/kg or 60 mg/kg of the KHK inhibitor or methylcellulose vehicle 1h before a fructose challenge test using 6 mg/kg of fructose or water and repeated blood samples were collected over time. The mice gavaged fructose and methylcellulose had elevated serum fructose excursion over 60 min, compared to the vehicle or inhibitor alone groups (**Fig S3F**). All three concentrations of the inhibitor (10, 30 and 60 mg/kg) showed a dose dependent increase in plasma fructose levels (**Fig S3F**). Serum glucose (**Fig S3G**) and insulin (**Fig S3H**) were the highest in mice treated with fructose and vehicle and they decreased with administration of 10, or 30 mg/kg of the inhibitor. Next, the mice were gavaged 10, or 30 mg/kg of the inhibitor and the level of the inhibitor in the blood was quantified over the 24h period. Based on *in vitro* potency 1 μM concertation of the inhibitor is required to inhibit KHK activity by 90% (IC90). Following 10 mg/kg administration of the inhibitor, *in vivo* concentration of the inhibitor above IC90 was sustained for seven hours (**Fig 3L**). Based on these studies, the mice in our experiment were gavaged 15 mg/kg of the inhibitor twice daily for four weeks. Exposure of the inhibitor was measured by mass spectrometry in the final plasma sample. The pharmacokinetic model fitting profile (**Fig 3M**) shows the free maximum exposure of the inhibitor was 4.6 μM and the free average exposure was 1.6 μM in line with *in vitro* IC90 reported in (Gutierrez et al., 2021). Therefore, the plasma concentration of the inhibitor achieved in this study was adequate to inhibit KHK activity, as previously published (Gutierrez et al., 2021).

### KHK siRNA treatment completely deletes KHK-C and increases HK2, while the inhibitor partially decreases both KHK-C and TKFC proteins

Given the differences in hepatic fructose metabolism, we next interrogated the fructose metabolism pathway (**Fig 4A**). The expression of enzymes catalyzing the first, second and third steps of fructose metabolism was increased 1.3-to 2.2-fold in mice on a HFD compared to LFD (**Fig 4B**). KD of KHK decreased both Khk-a and Khk-c mRNA better than 90% compared to the HFD group. Interestingly, inhibition of KHK activity also decreased Khk-a and Khk-c mRNA by 45-58% compared to the HFD group. KHK KD or inhibition of its activity did not affect aldolase b (Aldob) expression. The expression of the enzymes mediating the third step was not affected by KHK KD, except aldehyde dehydrogenase (Aldh3a2), which was 63% higher. Conversely, inhibition of KHK activity decreased triokinase and FMN cyclase (Tkfc) expression by 40% and it increased alcohol dehydrogenase (Adh1) 1.4-fold and Aldh3a2 3.2-fold, compared to the HFD group. Similar to mRNA, KHK-C protein was increased in mice on a HFD compared to a LFD (**Fig 4C, 4D**). The KHK+siRNA achieved a complete deletion of KHK-C protein, whereas, interestingly, the KHK+Inhib group had 56% lower KHK-C protein compared to the HFD group. ALDOB was slightly lower in both KHK+siRNA and KHK+Inhib groups. TKFC, which catalyzes a major arm of the third step of fructose metabolism was significantly lower in KHK+Inhib, but not in KHK+siRNA, compared to the HFD group. ADH1 was not different, whereas ALDH3 increased in both KHK+siRNA and KHK+Inhib groups. Together, these data indicate that KHK siRNA profoundly decreased KHK-C mRNA and protein, while KHK inhibitor decreased mRNA and protein of both KHK-C and TKFC.

**Figure 4.**
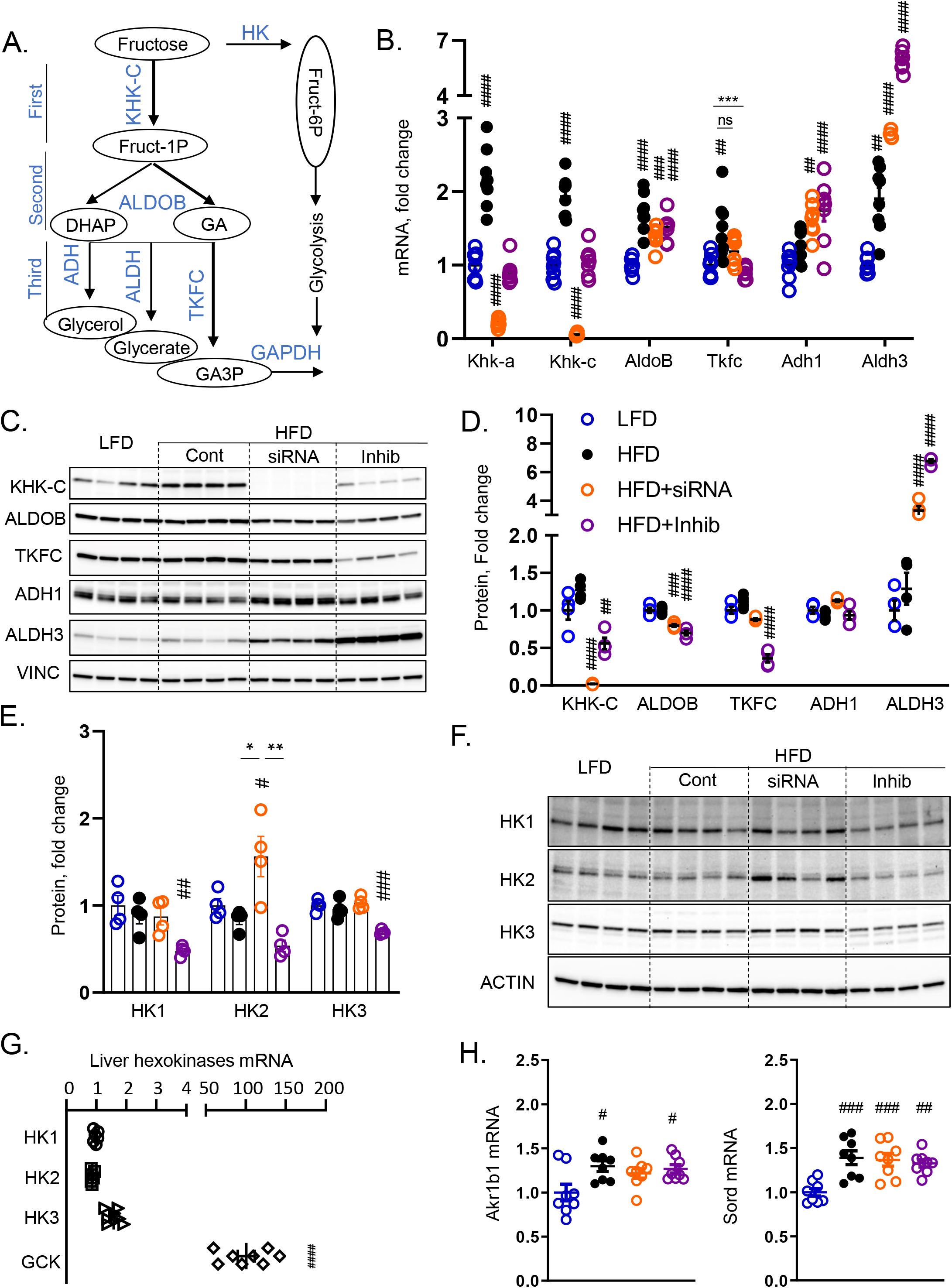
KHK siRNA completely deletes KHK-C and increases HK2, while the inhibitor partially decreases both KHK-C and TKFC proteins. (A) Fructose metabolism pathway. (B) mRNA of fructose metabolizing enzymes from livers of the mice. n=6 mice per group. (C) Western blot and (D) densitometry quantification of fructose metabolizing enzymes in liver lysates. n=4 mice per group. (E) densitometry quantification and (F) western blot of hexokinases in the liver. n=4 mice per group. (G) mRNA expression of hexokinases in the liver from mice fed a LFD. (H) hepatic mRNA expression of aldo-keto reductase (Akr1b1) and sorbitol dehydrogenase (Sord). Statistical analysis was performed using one way ANOVA compared to LFD group (#p < 0.05; ## p < 0.01; ### p < 0.001; #### p < 0.0001) with post hoc t tests between the individual groups (***p < 0.001).

In the absence of KHK, fructose may be metabolized by an alternative pathway where hexokinase (HK) phosphorylates fructose to fructose-6 phosphate (F6P). There are four hexokinases, one, two, three and four, a.k.a. Gck. All of these enzymes had similar mRNA expression in HFD+siRNA and HFD+Inhib groups, compared to the HFD, except HK2, which was the highest in the HFD+siRNA group (**Figs S4, 1J)**. Similarly, HK2 protein was only increased in the HFD+siRNA, but not KHK+Inhib group, compared to the HFD group (**Fig 4E, F**). The expression of hexokinases is much lower than Gck in the liver (**Fig 4G**). However, fasted fructose concentration (0.01 mM) (Kawasaki et al., 2002) is also much lower than glucose (5 mM), so low abundance of HK2 may still play a physiologic role in fructose metabolism. An increase in all three enzymes of fructose metabolism in mice on a HFD can be explained, in part, by endogenous fructose production mediated by aldo-keto reductase (Akr1b1) and sorbitol dehydrogenase (Sord) (**Fig 4H**). The endogenous fructose production pathway was not affected by KHK KD or its inhibition. In summary, KHK KD more completely abolished the fructose metabolism pathway leading to alternative metabolism via HK2. Conversely, the KHK inhibitor partially lowered both KHK-C and TKFC proteins, which may account for increased F1P in this group.

### RNAseq analysis reveals profound and unique roles of ketohexokinase knockdown versus inhibition on hepatic transcriptome

Given the unique difference in mRNA expression of the enzymes mediating fructose metabolism, we assessed global gene expression in these mice. The principal component analysis revealed that gene expression in the HFD group clustered differently than in LFD group (**Fig 5A**). Following KD of KHK, the gene expression clustered closer to the LFD group, while inhibition of KHK activity dramatically altered hepatic gene expression in a new direction. A heatmap analysis of top 40 genes identified a unique pattern of gene expression between the LFD and HFD groups, but also between KHK+siRNA and KHK+Inhib groups (**Fig 5B**). Some of the genes that were upregulated in HFD and decreased by both KHK KD and inhibition are involved in the regulation of cholesterol, such as Apoe4 and formation of atherosclerotic plaques, such as matrix metalloproteinase 12 (Mmp12). The genes that were decreased in HFD and restored by both KHK siRNA and inhibitor treatment are involved in ribosomal function, such as ribosomal protein S3 (Rps3) and ribosomal protein lateral stalk subunit P0 (RPLP0). The genes that were lowered by KHK KD, but not by the inhibitor treatment regulate lipid assemble, such as lysophosphatidylglycerol acyltransferase 1 (Lgpat1) and glycerol kinase (Gk). On the contrary, the genes that were normalized by the inhibitor, but not by KHK siRNA treatment were members of proteoglycans, such as glypican 1 (Gpc1) and proteoglycan 4 (Prg4). Interestingly, a number of genes regulating FAO were increased only in the inhibitor group, such as acyl-CoA thioesterase 2 (Acot2) and acyl-CoA oxidase 1 (Acox1). Next, we honed in on the unique pathways that were regulated exclusively by KHK KD or inhibition. The expression of the DNL pathway, regulated by transcription factor carbohydrate-responsive element-binding protein (ChREBP), was profoundly downregulated by KHK siRNA treatment (**Fig 5C**). On the other hand, KHK inhibition lead to upregulation of the peroxisomal FAO pathway, regulated by the peroxisome proliferator-activated receptor alpha (PPARα) transcription factor (**Fig 5D**).

**Figure 5.**
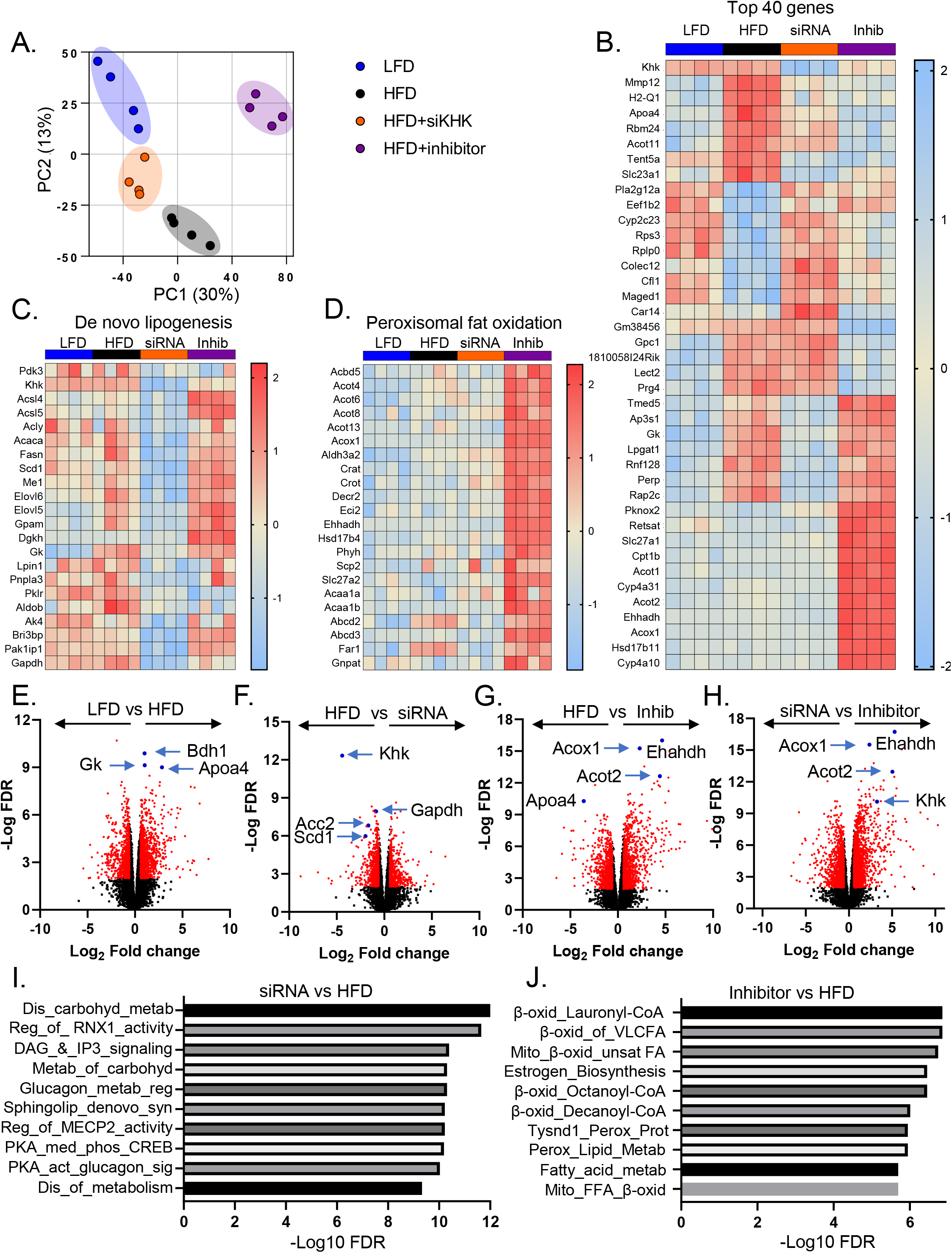
RNAseq analysis revealed profound and unique roles of ketohexokinase knockdown versus inhibition on hepatic transcriptome. (A) Principal component analysis or RNAseq data from RNA extracted from the liver of experimental mice (B) Heatmap of top 40 genes. (C) Heatmap of de novo lipogenesis pathway and (D) peroxisomal fatty acid oxidation genes. Volcano plot comparison of (E) HFD compared to LFD, (F) HFD compared to HFD+siRNA, (G) HFD compared to HFD+Inhibitor, and (H) HFD+siRNA compared to HFD+Inhibitor. Reactome pathway analysis showing the most significantly altered pathways between HFD+siRNA versus HFD group and (J) HFD+Inhibitor versus HFD group.

Volcano plot comparison of the individual groups revealed that the upregulated genes in HFD, compared to LFD group are involved in lipid homeostasis, including Gk, Apoa4 and 3-hydroxybutyrate dehydrogenase 1 (Bdh1) (**Fig 5E**). The most down regulated genes in the KHK siRNA group, compared to the HFD group, included the genes involved in fructose metabolism, Khk and Gapdh, but also the genes involved in fatty acid synthesis, such as stearoyl-CoA desaturase 1 (Scd1), and acetyl-CoA carboxylase (Acc2) (**Fig 5F**). The genes increased with the inhibitor treatment, compared to the HFD group, are involved in lipid oxidation, including Acot2, Acox1 and enoyl-CoA hydratase and 3-hydroxyacyl CoA dehydrogenase (Ehahdh) (**Fig 5G**). These same genes were upregulated in the KHK+Inhib group compared to the HFD+siRNA group, as these genes were uniquely upregulated by the inhibitor (**Fig 5H**). Reactome pathway analysis revealed that some of the most downregulated pathways in HFD+siRNA group mediate carbohydrate metabolism and de novo synthesis of lipids, such as sphingolipids (**Fig 5I**). Conversely, some of the most upregulated pathways in the HFD+Inhib group were mediating mitochondrial and peroxisomal FAO (**Fig 5J**). These changes persisted when the HFD+Inhib group was compared to the HFD+siRNA group (**Fig S5A**), since an increase in these genes was strongly driven by the inhibitor treatment.

An increase in FAO pathway and PPARα target genes with the inhibitor treatment was unexpected, so we assessed PPARα expression, which was elevated in the HFD and KHK+siRNA groups and was stepwise higher in HFD+Inhib group (**Fig S5B**). Next, we tested whether the inhibitor or F1P, produced by the inhibitor treatment, can bind to PPARα ligand binding domain to increase PPARα transcriptional activity in COS-7 cells transfected with PPARα luciferase reporter. As expected, WY14643, a PPARα agonist increased luciferase activity in a dose dependent fashion (**Fig S5C**), however, neither the inhibitor (**Fig S5D**) or F1P (**Fig S5E**) increased PPARα luciferase activity. Conversely, the cells treated with 25 μM of WY14643 and increasing doses of F1P showed slight, but significant reduction in PPARα activity following F1P treatment with concentrations above 100 μM (**Fig S5F, G**). Together, these data suggest that KHK KD decreased the DNL pathway, while the inhibitor treatment increased the FAO pathway mediated by PPARα. Furthermore, the inhibitor did not directly upregulate PPARα activity, whereas F1P may be a weak PPARα antagonist.

### Ketohexokinase knockdown decreased DNL, while the inhibitor increases FAO pathway

We verified some of the most interesting changes in RNAseq data by qPCR and Western blot. ChREBP-β (MLXIPL-β), the active isoform of the ChREBP transcription factor, was decreased in all mice on HFD, compared to the LFD group, since the HFD contains a reduced amount of carbohydrates (**Fig 6A**). MLXIPL-β, but not MLXIPL-α (**Fig S6A**), was further reduced with the siRNA or the inhibitor treatment, but the reduction was significantly greater in the HFD+siRNA group. The expression of sterol regulatory element-binding transcription factor 1c (SREBF1c) was elevated two-fold in all mice on HFD, compared to the LFD group, but there was no further effect with siRNA or the inhibitor treatment (**Fig 6B**). SREBF1a followed a similar expression pattern (**Fig S6B**). Nuclear translocation of ChREBP was similar in mice on HFD and LFD (**Fig 6C**). KHK siRNA decreased ChREBP compared to the HFD group. The inhibitor treatment did not significantly lower ChREBP. Nuclear translocation of SREBP1 was higher in mice on a HFD, compared to LFD group (**Fig 6C**). SREBP1 was not markedly affected in the HFD+siRNA and HFD+Inhib groups. The expression of downstream genes mediating lipogenesis was not elevated in mice on a HFD, and they were profoundly reduced in the HFD+siRNA, but not in the HFD+Inhib group (**Fig 6D**), consistent with the changes in ChREBP. The protein levels of these lipogenic enzymes were likewise profoundly decreased only in the HFD+siRNA group (**Figs 6E, S6C**), consistent with RNAseq data.

**Figure 6.**
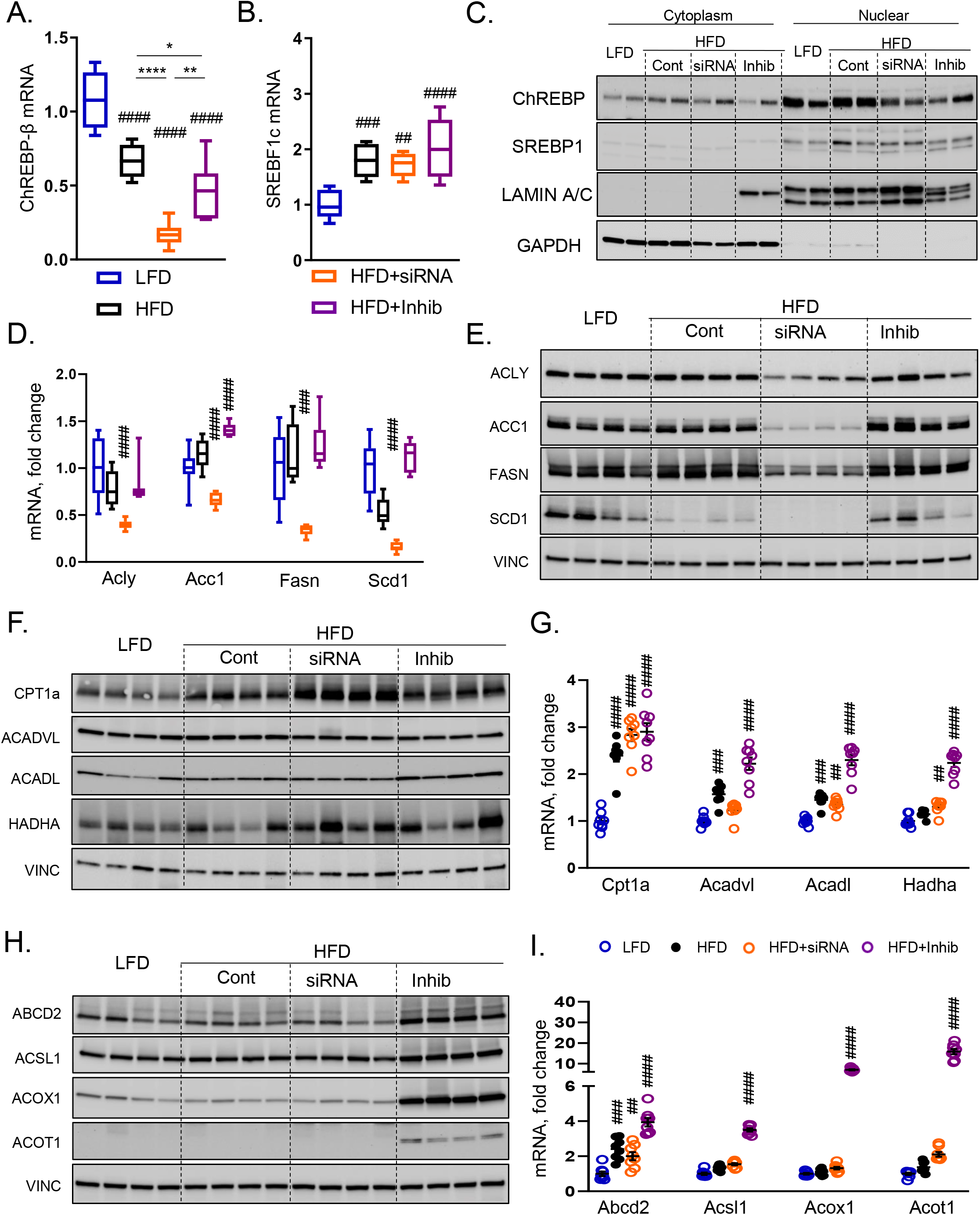
Ketohexokinase knockdown decreased DNL, while the inhibitor increases FAO pathway. (A) ChREBP-β (MLXIPL-β) and (B) SREBF1c mRNA expression in the liver of the experimental mice. (C) Western blot showing nuclear translocation of ChREBP, SREBP1 proteins in the liver. (D) mRNA and (E) Western blot quantification of proteins involved in de novo lipogenesis. (F) Protein and (G) mRNA expression of genes regulating mitochondrial fatty acid oxidation. (H) Protein and (I) mRNA expression of genes regulating peroxisomal fatty acid oxidation. n=7-8 mice per group for mRNA expression and n=4 mice per group for protein quantification. Statistical analysis was performed using one way ANOVA compared to LFD group (## p < 0.01; ### p < 0.001; #### p < 0.0001) with post hoc t tests between the individual groups (*p < 0.05; **p < 0.01; ****p < 0.0001).

Next, we quantified the protein levels of the enzymes mediating mitochondrial beta oxidation. CPT1α, the rate limiting enzyme of mitochondrial FAO was increased in HFD group compared to the LFD group (**Figs 6F, S6D**). CPT1α was further elevated in the HFD+siRNA, but not HFD+Inhib group. ACADVL and HADHA were not changed and ACADL was only increased in the HFD+Inhib group. In spite of the modest changes on protein level, mRNA expression of these genes were significantly elevated in mice on a HFD and they further increased in the HFD+Inhib, but not in the HFD+siRNA group (**Fig 6G**). The protein levels of enzymes mediating peroxisomal FOA were markedly elevated 2 to 240-fold only in the HFD+Inhib group (**Figs 6H, S6E**). Similarly, their mRNA expression was 4 to 20-fold higher in the HFD+Inhib group compared to the LFD group (**Fig 6I**). In summary, KHK KD markedly decreased DNL pathway, whereas KHK inhibitor profoundly increased peroxisomal FAO, in agreement with RNAseq data.

### In vitro overexpression of wild-type KHK supports glycogen accumulation, while overexpression of kinase dead mutant KHK increases expression of the DNL pathway

The differences between KHK KD and inhibition of its kinase activity suggest that there may be kinase independent functions of KHK. To test this hypothesis, we overexpressed GFP-tagged wild type mouse KHK-C (WT) or mouse kinase dead mutant KHK-C (KM) in human HepG2 control cells (CC) using lentivirus. KM KHK-C was generated by introducing a point mutation (G527R) in the ATP binding domain, so that the enzyme can bind fructose, but cannot phosphorylate it to F1P. For comparison, the KHK inhibitor also targets the ATP binding domain, similarly producing a nonfunctional enzyme. Compared to the control human cells, overexpression of WT or KM robustly induced mouse KHK-C expression (**Fig 7A**). We selected the clones with similar KHK-C overexpression for the subsequent analysis. KHK-C protein was much higher in the cells overexpressing WT, compared to KM suggesting decreased protein stability (**Fig 7B**). Indeed, when treated with cycloheximide, a protein synthesis inhibitor, KM KHK-C protein degraded faster than WT KHK-C protein (**Fig S7A**). Control HepG2 cells do not endogenously express KHK-C at high levels, but they also do not express ALDOB and TKFC needed for downstream fructose metabolism (**Figs 7B, S7B**). KHK enzymatic activity was markedly elevated in WT KHK-C cells, but not in KM KHK-C cells treated with fructose, confirming that kinase dead mutant enzyme is indeed not able to metabolize fructose (**Fig 7C**). Increased KHK activity in cells that do not express ALDOB or TKFC is expected to lead to the accumulation of F1P. F1P at high levels is known to be hepatotoxic. Indeed, WT KHK-C cells treated with 5 mM fructose, but not with 3-O methylfructose (3-OMF), a non-metabolizable form of fructose, exhibit higher cell injury leading to a lower total protein (**Fig 7D**). A decrease in total protein was not observed in control cells or KM KHK-C cells treated with fructose or 3-OMF. Hepatocyte injury was also measured by quantifying alanine aminotransferase (ALT) in cell supernatant. ALT was only elevated with fructose, but not glucose, treatment of WT KHK-C cells (**Fig 7E**). Thus, increased KHK activity in the cells treated with fructose that do not express downstream fructose metabolizing enzymes resulted in cell injury, likely due to F1P accumulation.

**Figure 7.**
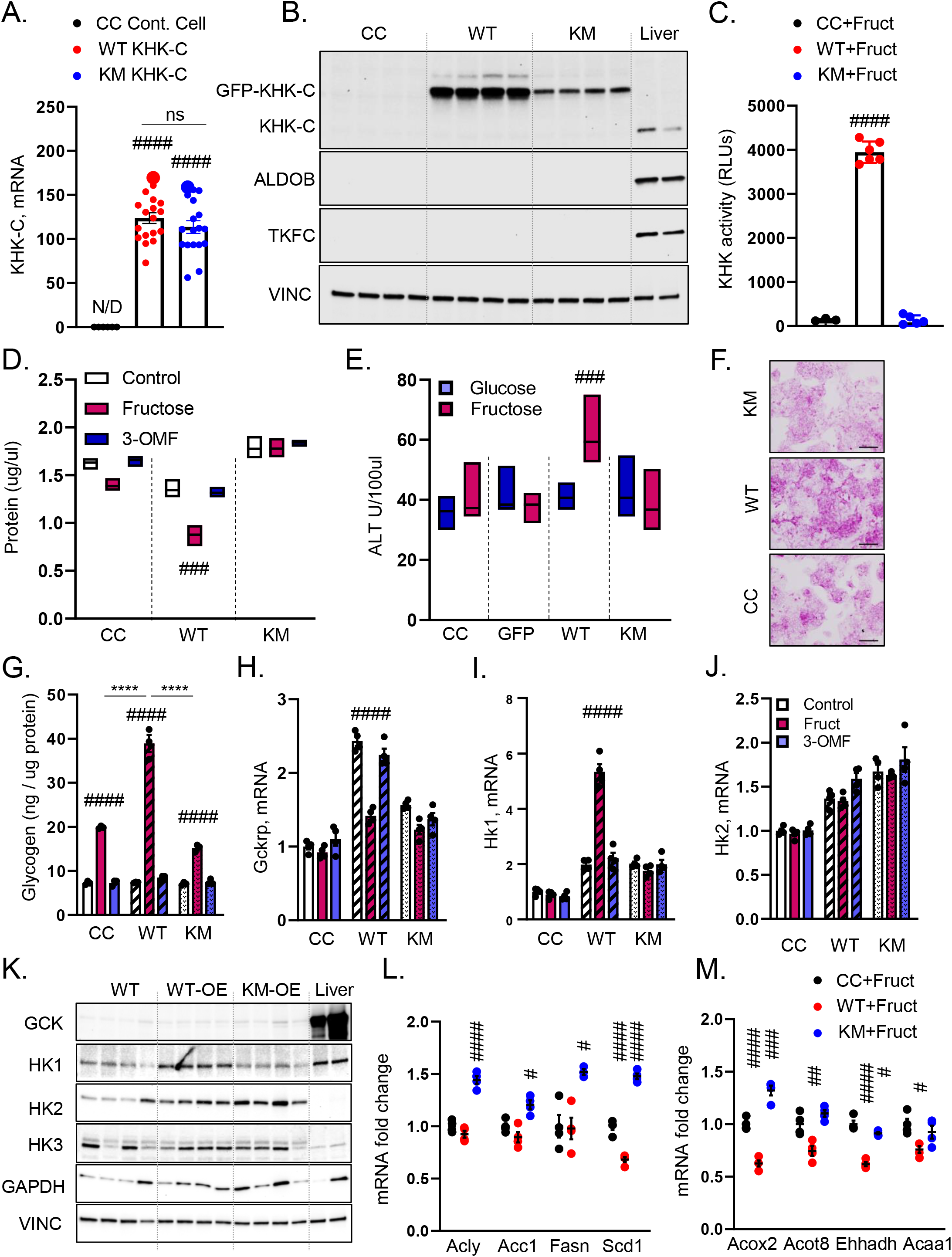
Overexpression of wild-type KHK supports glycogen accumulation, while overexpression of kinase dead mutant KHK increases expression of the DNL pathway. (A) KHK-C mRNA expression in control HepG2 cells, GFP-tagged wild type mouse KHK-C overexpressed cells (WT KHK-C), or mouse kinase dead mutant overexpressed (KM KHK-C) cells using lentivirus transfection. (B) protein levels of fructose metabolism related enzymes in control HepG2 cells, WT KHK-C cells, KM KHK-C cells, and mouse liver. (C) KHK activity in control HepG2 cells, WT KHK-C cells and KM KHK-C cells. (D) Total protein amount and (E) ALT level after treatment with 5mM fructose or 5mM 3-O methylfructose (3-OMF) for 24 hr. (F) Periodic acid-Schiff (PAS) staining of HepG2 cells treated with fructose for 24 hr. (G) glycogen quantification in control HepG2 cells, WT KHK-C cells and KM KHK-C cells treated with fructose or 3-OMF. mRNA of Gckrp(H), Hk1(I), and HK2(J) expression in control HepG2 cells, WT KHK-C cells and KM KHK-C cells treated with fructose or 3-OMF. (K) Western blot quantification of enzymes involved in sugar metabolism in control HepG2 cells, WT KHK-C cells, KM KHK-C cells, and mouse liver. (L) mRNA expression of de novo lipogenesis genes and (M) fatty acid oxidation genes. n=4 for mice per group for gene expression and protein quantification. Statistical analysis was performed using one way ANOVA compared to LFD group (#p < 0.05; ## p < 0.01; ### p < 0.001; #### p < 0.0001) with post hoc t tests between the individual groups (****p < 0.0001).

Similar to *in vivo* findings, glycogen deposition, as assessed by PAS staining, was increased in WT but not KM or CC cells treated with fructose (**Fig 7F**). We quantified intracellular glycogen, which was increased with fructose compared to glucose treatment in all cells. Glycogen was highest in the cells overexpressing WT KHK-C (38.9±2.0ng/ug), compared to CC (19.9±0.2ng/ug) or KM (15.2±0.5ng/ug) cells treated with fructose or 3-OMF (**Fig 7G**). Fructose supports glycogen accumulation by enhancing glucokinase activity (Vandercammen et al., 1992). Gck was not expressed in HepG2 cells, but Gckrp was increased over 2-fold in WT KHK-C cells and it completely normalized following fructose, but not 3-OMF treatment (**Fig 7H**). In the absence of Gck, Gckrp may inhibit hexokinase activity and expression. Indeed, Hk1 expression was 5-fold higher in WT KHK-C cells treated with fructose (**Fig 7I**). Hk2 expression was elevated in both WT and KM KHK-C cells, but it was not affected by fructose or 3-OMF treatment (**Fig 7J**). In agreement with mRNA, GCK protein was not abundant in HepG2 cells compared to mouse liver (**Fig 7K, S7C**). HK1 protein was only elevated in WT KHK-C cells, whereas HK-2 was increased 2-3 fold in both WT and KM cells, compared to CC (**Fig 7K, S7D**). HK3 was not significantly altered, while GAPDH tended to be higher in both WT and KM cell.

Lastly, we interrogated DNL and FAO pathways in these cells treated with 5 mM fructose. Fructose-fed WT KHK-C overexpressing cells did not show an increase in Acly, Acc1, Fasn and Scd1 expression (**Fig 7L**). Interestingly, the cells overexpressing KM KHK-C had significantly higher expression of DNL enzymes compared to CC suggesting that the DNL pathway is regulated, in part, through kinase independent function of KHK-C. This correlated with increased expression of ChREBF and SREBP1c lipogenic transcription factors in KM but not WT cells (**Fig S7E, F**). On the other hand, fructose fed WT KHK-C cells, but not KM KHK-C cells had decreased expression of genes regulating FAO (**Fig 7M**). These data indicate that fructose metabolism and thus KHK activity is required to decrease FAO observed with fructose intake. In summary, KHK activity is required to generate F1P, induce glycogen accumulation and decrease FAO, whereas DNL pathway is increased independent of KHK-C kinase function.

## DISCUSSION

In the present study, we have compared the effects of liver-specific knockdown of KHK versus systemic inhibition of KHK kinase activity on hepatic metabolism. We find that these two modalities targeting the same enzyme exert uniquely different metabolic outcomes. KHK siRNA treatment completely and specifically deletes KHK protein in the liver. This leads to the upregulation of an alternative pathway of fructose metabolism via HK2, which converts fructose to F6P, an intermediate in glycolysis. An increase in the glycolysis pathway following KHK KD likely accounts for improved glucose tolerance in these mice. However, KHK inhibitor decreases both KHK and TKFC proteins, resulting in F1P accumulation in the liver. Elevated F1P is associated with glycogen accumulation, hepatomegaly and impaired glucose tolerance. Perhaps the most striking difference between KHK KD and inhibition of its kinase activity is in the regulation of hepatic gene expression. KHK KD profoundly decreases expression of genes in the DNL pathway, which is not observed following the inhibition of its kinase activity. Similarly, overexpression of kinase dead mutant KHK-C increases the expression of DNL genes suggesting kinase independent role of KHK. On the other hand, the KHK inhibitor uniquely upregulates the expression of PPARα target genes mediating FAO. These effects are not directly facilitated by the inhibitor or F1P, while fructose metabolism *in vitro* lowers the expression of FAO genes. In summary, liver-specific KHK KD exerts beneficial effects on hepatic metabolism, while systemic inhibition of its kinase activity increases hepatic glycogen accumulation via its off targets effects on TKFC.

Liver-specific KHK KD induces a more complete abrogation of fructose metabolism as evident by minimal KHK activity, reduced hepatic F1P/Fructose ratio, and activation of an alternative pathway of fructose metabolism via HK2. In spite of achieving adequate plasma IC90 concentration, fructose metabolism in the liver is only partially reduced by the inhibitor as indicated by incomplete reduction of KHK activity, normal F1P/Fructose ratio and no improvement in systemic glucose tolerance. The F1P/Fructose ratio is higher in the inhibitor treated mice than expected given that partial inhibition of KHK activity should reduce fructose metabolism and thus F1P levels. Instead, F1P concentration was the highest in the HFD+Inhih group, likely due to a lower TKFC protein leading to a decrease in the third step of fructose metabolism. A decrease in KHK and TKFC is not entirely explained by a decrease in ChREBP, which is actually lower following KHK KD than in the inhibitor treated mice. Lower KHK and TKFC may be due to the inhibitor targeting the ATP binding domain of both KHK and TKFC, leading to protein destabilization. KHK and TKFC are the only kinases within the first three steps of fructose metabolism that possess ATP binding domain. Indeed, inhibiting ATP binding domain *in vivo* or introducing a point mutation in ATP binding domain *in vitro* lowers KHK protein. While both KHK and TKFC kinases contain ATP binding domain we do not have direct evidence that the KHK inhibitor can bind TKFC.

F1P is a toxic metabolite that leads to hepatic glycogen accumulation and hepatocellular injury (Singh and Sarma, 2022). Cytotoxic properties of F1P have been documented from yeast (Gibney et al., 2018) to human cells (Phillips et al., 1968). We observe lower total protein and higher ALT in fructose treated cells overexpressing KHK. We also see increased glycogen accumulation in the livers of mice with elevated F1P and in cells that can metabolize fructose. Glycogen accumulation can be accounted largely by an increase in F1P. Glucokinase is bound by glucokinase regulatory protein that inhibits its activity (Veiga-da-Cunha and Van Schaftingen, 2002). F1P promotes the dissociation of the complex and allows the production of substrate for glycogenesis (Vandercammen et al., 1992). Additionally, F1P increases the activity of glycogen synthase (Gergely et al., 1985). Another explanation for higher glycogen accumulation is a decrease in serum glucagon. These processes are interrelated as glucagon plays a role in regulating F1P levels (Davies et al., 1990). The strong propensity of F1P to stimulate glycogen accumulation is observed in the cells that overexpress wild type, but not kinase dead KHK. These cells also lack ALDOB required for further metabolism of F1P. *In vitro*, fructose metabolism decreases Gckrp expression and leads to upregulation of hexokinase 1 that may provide substrate for glycogen synthesis. Indeed, a dual role of F1P to support glycogenesis and induce hepatotoxicity is best observed in patients with hereditary fructose intolerance (HFI) due to ALDOB deficiency (Singh and Sarma, 2022). The symptoms of HFI can be completely avoided in mice by knocking out KHK in order to prevent fructose metabolism to F1P (Lanaspa et al., 2018). Interestingly, in our study an increase in F1P following the inhibitor treatment induced glycogen accumulation, but was not associated with increased liver injury or inflammation.

Both KD of KHK and inhibition of its kinase activity effectively resolve hepatic steatosis. However, these modalities reduce hepatic fat accumulation via substantially different mechanisms. RNAseq analysis shows that a complete absence of KHK protein, but not inhibition of its kinase activity uniquely decreases the expression of genes mediating DNL. This is confirmed by markedly lower mRNA and protein levels of DNL enzymes, such as ACLY, ACC1, FASN and SCD1 in the HFD+siRNA, but not the HFD+Inhib group. The effects on DNL are likely mediated by kinase independent effects of KHK, as overexpression of kinase dead mutant KHK was sufficient to increase mRNA of DNL genes. We recently reported the first evidence pointing to kinase independent effects of KHK in mediating endoplasmic reticulum (ER) stress (Park et al., 2023). Our subsequent work suggests that kinase independent function of KHK mediates acetylation of proteins regulating hepatic metabolism (Helsley et al., 2023). Acetylated proteins may not fold properly, triggering misfolded protein response and ER stress. Alternatively, the effects of KHK on DNL may be, in part, mediated by a greater reduction in ChREBP. Indeed, acetylation of histones is a powerful mode of regulating gene expression and acetylation of ChREBP increases its activity (Bricambert et al., 2010). The KHK inhibitor has been reported to reduce DNL in rats (Gutierrez et al., 2021). We may not have observed lower DNL as we did not use a high-fructose diet, we used a lower dose of the inhibitor while only 30 mg/kg dose was reported to reduce DNL and mice are less sensitive to the DNL effects of dietary fructose than rats. On the other hand, the small molecule inhibitor uniquely and profoundly increases the expression of genes regulating FAO. This is not mediated by direct binding of the inhibitor to PPARα ligand binding domain. However, the inhibitor could stimulate some of the PPARα co-activators, perhaps PGC1α, and thus indirectly enhance PPARα transcriptional activity. Another possibility for enhanced FAO pathway is a decrease in TKFC. A knockout of TKFC in the liver shifts the third step of fructose metabolism toward the ALDH pathway resulting in increased FAO (Liu et al., 2020). In our study, ALDH was indeed more strongly upregulated in the HFD+Inhib, than in the HFD+siRNA group. The inhibitor-induced decrease in TKFC and strong upregulation of the PPARα pathway may explain lower steatosis without a substantial decrease in the DNL pathway.

Fructose intake is a well-recognized risk factor for the development of obesity and metabolic dysfunction. We show that targeting KHK via liver-specific siRNA completely and specifically prevents fructose metabolism and results in the resolution of hepatic steatosis and improves glucose tolerance. On the other hand, small molecule inhibitor of KHK only partially reduces KHK activity in the liver, but it also targets TKFC resulting in elevated F1P, glycogen accumulation, hepatomegaly and no improvement in glucose tolerance. Our study reaffirms that targeting KHK is still a viable option for the management of NALFD. However, methods, such as antisense oligonucleotides, GalNAc-siRNA or new crisper Cas11 technologies that induce a complete deletion of KHK protein, thus abolishing the enzyme’s kinase dependent and independent functions, hold greater promise to completely reverse metabolic dysfunction induced by dietary fructose.

## LIMITATIONS OF THE STUDY

KHK inhibitor concentration above 1μM was sustained for seven hours following a single dose administration. We gavaged mice twice daily, which was stressful enough to prevent further weight gain, so that three times per day gavage was not feasible. This means that the mice had ten hours per day of uninterrupted fructose metabolism. However, a more consistent inhibition of KHK is expected to lead to higher F1P levels and even stronger glycogen accumulation and liver injury.

## AUTHOR CONTRIBUTIONS

Conceptualization, S.H.P., S.S.; Methodology, L.R.C., H.C, R.C.S., K.W., G.O., J.B., T.D.H.; Formal Analysis, L.R.C., R.C.S.; Investigation, S.H.P., T.F., L.R.C., H.C., R.C.S., K.W., J.B. G.O., A.B., M.P., T.D.H., G.J.M.; Resources, K.W., G.O., J.B., S.D.; Data Curation, L.R.C., R.C.S.; Writing – Original Draft, S.H.P., S.S.; Supervision, S.S.; Project Administration, S.H.P., T.F., S.S.; and Funding Acquisition, S.S.

## ACKNOWLEDGEMENT

The authors would like to thank Leila Noetzli, Ho-Chou Tu, and Kevin Fitzgerald at Alnylam Pharmaceutical for their assistance with the project and for providing KHK siRNA. Additionally, we would like to thank Mark Keibler for quantifying F1P using mass spectrometry at Alnylam Pharmaceutical. We appreciate Terry Flier and Anna Borodovsky at Alnylam Pharmaceutical for critically reading the manuscript. Next, we would like to thank Jian-Ming Liu for measuring the *in vitro* potency of the inhibitor and Sara Lindblom for formulating the inhibitor for oral administration at AstraZeneca. We thank Gregory Graft for providing ABCD2 antibody. We thank the BPF Genomics Core Facility at Harvard Medical School for their expertise and instrument availability that supported RNAseq work.

## TABLES WITH TITLES AND LEGENDS

Supplemental Table 1. Real-time quantitative mouse PCR primers and antibodies used in the study.

Supplemental Table 2. Real-time quantitative human PCR primers.

Supplemental Table 3: Antibodies used for Western blot

## RESOURCE AVAILABILITY

### Lead contact

Additional information and requests for resources and reagents should be directed to and fulfilled by the Lead Contact, Samir Softic.

### Materials availability

All unique/stable reagents generated in this study are available from the Lead Contact with a completed Materials Transfer Agreement.

### Data and code availability

Will deposit RNA seq data to GEO, a public archive, once the paper is accepted.

## STAR METHODS

### Animals and Diets

All animal protocols were in accordance with NIH guidelines and approved by the IACUC of the University of Kentucky. Mice were housed at 20-22 LC on a 12 h light/dark cycle with *ad libitum* access to food and water. C57BL/6J male mice at 6 weeks of age were purchased from Jackson Laboratory and fed either low-fat diet (LFD, Research diets #12450K) or 60% high-fat diet (HFD, Research diets, D12492) for 6 weeks. Thereafter, the mice on high-fat diet were subdivided into three groups; control group (HFD), KHK siRNA injected group (HFD+siRNA) and KHK inhibitor gavaged group (HFD+Inhib). To ensure equal stress/treatment, all mice were injected with either luciferase control siRNA (10 mg/kg) or siRNA targeting total KHK (20 mg/kg) every two weeks and were gavaged either methylcellulose control or (15mg/kg) KHK inhibitor (PF-06835919) in methylcellulose, twice daily for four weeks. The animals were weighed, and their food intake was recorded once per week. GTT and ITT were performed after 8 and 9 weeks on the diets, respectively. Mice were sacrificed after 10 weeks on the diets. First, the mice were gavaged and then fasted for 2 hours. The mice were injected saline or 1U of insulin (Humulin R-Lilly #HI-213) via inferior vena cava 10 minutes before the tissue was collected in liquid nitrogen. One mouse from each cage, i.e., dietary group, was utilized before sacrificing the next mouse in the same cage. This was repeated until all four mice per cage were sacrificed. Necropsy started at 9 am and finished at 12 am. The time that each mouse was sacrificed was recorded to calculate the length of the exposure after the last inhibitor dose.

### Liver-specific KHK knockdown

Liver-specific knockdown was achieved by utilizing an siRNA conjugated to N-acetylgalactosamine (GalNAc). Alnylam Pharmaceuticals synthesized siRNA to specifically target mouse total Khk mRNA. The siRNA consists of 2 strands, guide and passenger. The guide strand carries the sequence information necessary for target-gene recognition, while the passenger strand supports loading into the RNA-induced silencing complex (RISC). siRNA has undergone chemical modifications to achieve long-lasting effect and specificity for hepatocytes. The combination of the 2′-fluoro, 2′-O-methyl and phosphorothioate modifications provides protection against exonuclease degradation, allowing for marked compound stabilization. The guide strand is conjugated to a trivalent GalNAc specifically recognized by the asialoglycoprotein receptor (ASGPR), which is highly expressed on the surface of hepatocytes, achieving hepatocyte-specific delivery and uptake. A GalNAc conjugate siRNA targeting luciferase was used as a negative control. Mice were injected subcutaneously with 20 mg/kg of KHK siRNA, or 10 mg/kg of luciferase control siRNA every two weeks.

### KHK inhibitor dose calculations and pharmacokinetic studies

KHK inhibitor, PF-06835919 was synthesized at Pharmaron based on the structure of described in (Futatsugi et al., 2020). The *in vitro* potency, 5 nM, of the cpd was confirmed using the method described in (Park et al., 2021). The compound was formulated in 0.5% HPMC 10000 cP, 0.1% Tween80, pH 9. The suspension was divided into 10 ml tubes for each dosing occasion and frozen. Before each dosing, the formulation was thawed, allowed to reach room temperature, and then stirred for at least 30 min. The dose of 15 mg/kg gavaged twice daily was based on PK modelling as described in the results and a pilot fructose tolerance test confirmed that the dose of the inhibitor used was adequate to inhibit KHK activity *in vivo*. Pilot fructose tolerance test: The experimental procedures were approved by the Gothenburg, Sweden ethics review committee on animal experiments (2002-2019). Male c57bl/6J mice on 60 % HFD (Research Diets D12492) starting at 6 weeks age, were purchased at 16 weeks of age from Taconic Denmark and kept in an Association for Assessment and Accreditation of Laboratory Animal Care (AAALAC) accredited facility with environmental control; 20–22°C, relative humidity 40–60%, a 12-hour light-dark cycle (lights off at 6 pm). At the time of the challenge, the mice weighed 49.9±0.2g. Prior to the fructose tolerance test, the mice were gavaged 10 mg/kg, 30 mg/kg or 60 mg/kg of the KHK inhibitor or methylcellulose vehicle. One hour later, 6 mg/kg of fructose or water was gavaged and repeated blood samples were collected over 1h. Fructose was analysed using BioAssay Systems EnzyChrom™ Fructose assay Kit (EFRU-100), glucose was analysed using a handheld ACCU-chek device and insulin was measured by ELISA (Chrystal Chem Ultra Sensitive Mouse Insulin ELISA Kit (Cat # 90080).

Plasma was collected for exposure analysis at the termination of the main experiment. The plasma samples were precipitated with 100% acetonitrile. After vortexing and centrifugation the supernatant was transferred to a new plate and diluted 1:2 with H_2_O, 0.2% formic acid. Matrix matched calibration samples, quality controls for PF-06835919 and blanks were prepared in the same way as the study samples. The study samples, calibration samples and quality controls were injected on an Acquity Ultra Performance LC coupled to a Sciex API 4500 mass spectrometer. The analytical column was a BEH C18, 1.7 µm, 2.1 x 50 mm kept at 60°C. The flow rate was 0.750 mL/min using a mobile phase of 2% acetonitrile and 0.2 % formic acid in water (A) and 0.2% formic acid in acetonitrile (B). A gradient run was applied where 35% B at 0.0 minutes was increased to 100% B at 1.2 to 1.5 minutes then 35% B was applied for conditioning from 1.5 to 1.8 minutes. The mass spectrometry method was operated in positive electrospray mode using multiple reaction monitoring of transitions 357.2 >315.3 for PF-068359198. The data from the mass spectrometer were processed where the samples were read of a linear fit curve based on the calibration samples. All CVs and biases were below 10 %. Due to the short half-life in preclinical species (Futatsugi et al., 2020) PF-06835919 was orally administered twice daily to ensure adequate exposure during the study. The predicted exposure associated with this dose level has previously shown high KHK inhibition in rats and human and has proved reversal of metabolic dysfunction induced by high fructose diet (Gutierrez et al., 2021). Plasma concentration of PF-06835919 was measured from 2 until 5 hours post last dose. Therefore we fitted these data with 1 compartmental pharmacokinetic model to assess if average plasma concentration was in line with predicted target exposure.

### Glucose and insulin tolerance test

For glucose tolerance test (GTT), mice were fasted overnight and injected intraperitoneally (IP) with 2g glucose per kg of body mass. Blood glucose levels were measured at 0, 15, 30, 60, and 120 minutes using a glucose meter (Infinity, US Diagnostics). Insulin tolerance tests (ITT) were performed in nonfasted mice by i.p. injection of 1 mU insulin per kg of body mass. Blood glucose levels were measured at indicated times.

### Liver Histology and serum assays

Histology sections were prepared from formalin fixed paraffin embedded liver sections. H&E staining and Periodic acid–Schiff staining were performed by the University of Kentucky Pathology Research Core. Liver homogenates were used for the quantification of triglycerides (Pointer scientific, T7532-1L) following manufacturer’s guidelines, as previously published (Softic et al., 2012). Plasma insulin was quantified using an ELISA kit (Crystal chem, 90080). ALT levels were measured using commercial ALT kit (Catachem, C164-0A). Urinary fructose was quantified using a fructose assay kit (BioAssay Systems, EFRU-100).

### PPAR**α** Transcriptional Activity Assay

We used Cos7 green kidney monkey cells that were routinely cultured and maintained in Dulbecco’s modified Eagle’s medium containing 10% fetal bovine serum (FBS) with 1% antibiotic-antimycotic. Cultures were maintained at 37 °C and a 5% CO_2_ saturation. We performed the assay as we have previously described in (Gordon et al., 2020). In brief, PPARα transcriptional activity was measured by transiently transfecting Cos7 cells using the Neon^TM^ Transfection System per the manufacturer’s settings, with a plasmid containing the 9x upstream activation sequence (9xUAS) GAL4-driven promoter luciferase vector construct (9x UAS-Luc, 16μg) along with a plasmid containing the GAL4 DNA-binding domain fused with the LBD of PPARα (GAL4-PPARα-LBD, 6μg), and a pRL-CMV plasmid for Renilla luciferase expression (0.25μg) as a transfection normalizing control, as we previously described in (Gordon et al., 2020). 24 hours after electroporation, warm DMEM + 10% dialyzed FBS + 1% AA was added to the cells. After an additional 24 hours, the cells were treated with a dose-dependent increase of WY14643 (0.125 - 50 μM), KHKi (0.125 - 10 μM), Fructos-1-Phosphate (100 nM - 1mM), and Fructos-1-Phosphate (100 nM - 1mM) + 25μM WY14643 for 24 h. The activity was then measured by dividing Firefly luciferase with the Renilla luciferase using the Dual-Luciferase^®^ reporter assay system (Promega) with the Promega GloMax^®^ 96 microplate luminometer (Promega).

### qPCR and mRNA quantification

Gene expression was quantified as previously described (Softic et al., 2016a). Briefly, mRNA was extracted by homogenizing liver tissue in trizol, treating with chloroform and participating in 70% ethanol. mRNA was purified using RNeasy Mini Kit columns (Qiagen, #74106). cDNA was made using High Capacity cDNA Revers Transcription Kit (Applied Biosystems, #4368813). qPCR was performed utilizing C1000 Thermal Cycler (BioRad, CFX384) and QuantStudio™ 7 Flex Real-Time PCR System (TermoFisher Scientific, 4485701). Mouse and human primer sequences used are listed in the supplemental tables 1 and 2, respectively. An average of 18S and TBP was used as housekeeping genes to normalize the mRNA data.

### Protein Extraction and Immunoblot

Tissues were homogenized in RIPA buffer (EMD Millipore) with protease and phosphatase inhibitor cocktail (Bimake.com, B14002, B15002). Proteins were separated using SDS-PAGE and transferred to PVDF membrane (Millipore). Immunoblotting was achieved using the indicated antibodies listed in the supplemental table 3. Images were captured by ChemiDoc MP imaging system (BioRad, 12003154) and iBright imaging system (Thermo Fisher, CL1000). Quantification of immunoblots was performed using ImageJ.

### Generation of Stable KHK-C Overexpressing HepG2 cells

Human hepatic HepG2 cells were cultured under standard conditions in 1:1 Dulbecco’s Modified Eagle Medium (DMEM) and Ham’s F-12 media (Corning) supplemented with 10% fetal bovine serum (FBS, Corning) and 1% penicillin-streptomycin in a 5% CO_2_-humidified cell culture incubator at 37°C. Lentiviral particles were purchased from Origene containing the mouse KHK-C sequence (MR204149L2V) or Kinase dead mutated KHK-C fused to GFP (custom ordered from Origene). HepG2 cells were transduced with lentivirus at a MOI of 5 and media was changed every two days. After 5 days, cells were sorted for GFP-intensity and only GFP+ cells were collected in 96-well plates and propagated for an additional 2-3 weeks. KHK-C was quantified in these clones by both QPCR and western blot. The highest expressing clones were used for subsequent experiments.

### Matrix-Assisted Laser Desorption/Ionization Mass Spectrometry Imaging of Glycogen In Situ

MALDI imaging of in situ glycogen was performed based on the previously described technique (Young et al., 2022). Formalin-fixed paraffin embedded (FFPE) tissues or purified glycogen were sectioned at 5 µm or spotted directly onto glass slides. The slide went through dewaxing and rehydration washes, followed by antigen retrieval in a citraconic anhydride buffer solution. An HTX M5 robotic sprayer was used to coat tissues in a solution of isoamylase. Following the application of enzyme, the slides were incubated for 2 hours at 37^0^C in a humidity chamber, then dried overnight in a vacuum desiccator. The following day, the robotic sprayer coated the slides with a 7 mg/mL CHCA matrix (0.040 g of α-cyano-4-hydroxycinnamic acid in 5.7 mL of 50% Acetonitrile/0.1% Trifluoroacetic Acid solution) and the slides were stored in a desiccator until MALDI-MSI analysis could be performed. The slides were loaded into a Waters Synapt G2 SX mass spectrometer that used a UV laser of 100 µm in size to detect N-glycans and glycogen. Data analysis was then performed using Waters High Definition Imaging (HDI) software.

### RNA Sequence Analysis and Bioinformatics Methods

HTG EdgeSeq mRNA sequence analysis was performed by the BioPolymers Facility at Harvard Medical School. Reads were aligned to the mouse transcriptome with Kallisto and the transcript counts were converted to gene counts with tximport. One sample was supposed to be KHK-treated, but its KHK expression was as large as the non-treated, and thus this sample was removed. Genes must have at least one count per million (CPM) in at least four samples, or they are filtered out. We normalized expression with trimmed mean of M-values (TMM) and transformed to logCPM with Voom. To discover the differential genes, we use limma, an R package that powers differential expression analyses. We applied the Fry function of the Rotation Gene Set Test (Roast) method in the limma R package to perform pathway analysis. We obtained the Gene sets from the MSigDB Collections. We selected the gene sets that belong to the canonical pathways (CP) and Gene Ontology (GO). To get an overall view of the similarity and/or difference of the samples, we performed principal component analysis (PCA). In the volcano plot, the top genes with either the smallest p-values or largest logFC are labeled. In the heatmap, the same set of top genes are used, and the z-scores of the logCPM are plotted. Pathway analysis was performed as in our previously published work (Moreau et al., 2022).

### Statistical Analyses

All data are presented as mean ± SEM. The data analysis, comparing the effects of control to experimental conditions was first analyzed using one-way analysis of variance (ANOVA) with Dunnett’s multiple comparisons test for comparison of the individual groups. Significant difference among control and experimental groups is noted with a number sign (#) and significant difference between the individual groups under a black line is designated by an asterisk (*). For both signs, the single symbol represents a p value of <0.05, two symbols represent a p value of <0.01, and three symbols denote a p value of <0.001, and four symbols stand for a p value of <0.0001 throughout the study.

## METHOD DETAILS

For a detailed description of materials and methods and their respective catalog numbers, please refer to the Key resources table.

## REFERENCES

Abdelmalek, M.F., Suzuki, A., Guy, C., Unalp-Arida, A., Colvin, R., Johnson, R.J., Diehl, A.M., and Nonalcoholic Steatohepatitis Clinical Research, N. (2010). Increased fructose consumption is associated with fibrosis severity in patients with nonalcoholic fatty liver disease. Hepatology 51, 1961–1971.

Andres-Hernando, A., Orlicky, D.J., Kuwabara, M., Ishimoto, T., Nakagawa, T., Johnson, R.J., and Lanaspa, M.A. (2020). Deletion of Fructokinase in the Liver or in the Intestine Reveals Differential Effects on Sugar-Induced Metabolic Dysfunction. Cell Metab 32, 117–127 e113.

Bricambert, J., Miranda, J., Benhamed, F., Girard, J., Postic, C., and Dentin, R. (2010). Salt-inducible kinase 2 links transcriptional coactivator p300 phosphorylation to the prevention of ChREBP-dependent hepatic steatosis in mice. J Clin Invest 120, 4316–4331.

Canadian Diabetes Association Clinical Practice Guidelines Expert, C., Dworatzek, P.D., Arcudi, K., Gougeon, R., Husein, N., Sievenpiper, J.L., and Williams, S.L. (2013). Nutrition therapy. Can J Diabetes 37 Suppl 1, S45–55.

Chiang Morales, M.D., Chang, C.Y., Le, V.L., Huang, I.T., Tsai, I.L., Shih, H.J., and Huang, C.J. (2022). High-Fructose/High-Fat Diet Downregulates the Hepatic Mitochondrial Oxidative Phosphorylation Pathway in Mice Compared with High-Fat Diet Alone. Cells 11.

Cohen, C.C., Li, K.W., Alazraki, A.L., Beysen, C., Carrier, C.A., Cleeton, R.L., Dandan, M., Figueroa, J., Knight-Scott, J., Knott, C.J., et al. (2021). Dietary sugar restriction reduces hepatic de novo lipogenesis in adolescent boys with fatty liver disease. J Clin Invest 131.

Conroy, L.R., Stanback, A.E., Young, L.E.A., Clarke, H.A., Austin, G.L., Liu, J., Allison, D.B., and Sun, R.C. (2021). In Situ Analysis of N-Linked Glycans as Potential Biomarkers of Clinical Course in Human Prostate Cancer. Mol Cancer Res 19, 1727–1738.

Davies, D.R., Detheux, M., and Van Schaftingen, E. (1990). Fructose 1-phosphate and the regulation of glucokinase activity in isolated hepatocytes. Eur J Biochem 192, 283–289.

European Association for the Study of the, L., European Association for the Study of, D., and European Association for the Study of, O. (2016). EASL-EASD-EASO Clinical Practice Guidelines for the Management of Non-Alcoholic Fatty Liver Disease. Obes Facts 9, 65–90.

Futatsugi, K., Smith, A.C., Tu, M., Raymer, B., Ahn, K., Coffey, S.B., Dowling, M.S., Fernando, D.P., Gutierrez, J.A., Huard, K., et al. (2020). Discovery of PF-06835919: A Potent Inhibitor of Ketohexokinase (KHK) for the Treatment of Metabolic Disorders Driven by the Overconsumption of Fructose. J Med Chem 63, 13546–13560.

Gergely, P., Toth, B., Farkas, I., and Bot, G. (1985). Effect of fructose 1-phosphate on the activation of liver glycogen synthase. Biochem J 232, 133–137.

Gibney, P.A., Schieler, A., Chen, J.C., Bacha-Hummel, J.M., Botstein, M., Volpe, M., Silverman, S.J., Xu, Y., Bennett, B.D., Rabinowitz, J.D., et al. (2018). Common and divergent features of galactose-1-phosphate and fructose-1-phosphate toxicity in yeast. Mol Biol Cell 29, 897–910.

Gordon, D.M., Neifer, K.L., Hamoud, A.A., Hawk, C.F., Nestor-Kalinoski, A.L., Miruzzi, S.A., Morran, M.P., Adeosun, S.O., Sarver, J.G., Erhardt, P.W., et al. (2020). Bilirubin remodels murine white adipose tissue by reshaping mitochondrial activity and the coregulator profile of peroxisome proliferator-activated receptor alpha. J Biol Chem 295, 9804–9822.

Gutierrez, J.A., Liu, W., Perez, S., Xing, G., Sonnenberg, G., Kou, K., Blatnik, M., Allen, R., Weng, Y., Vera, N.B., et al. (2021). Pharmacologic inhibition of ketohexokinase prevents fructose-induced metabolic dysfunction. Mol Metab 48, 101196.

Helsley, R.N., Moreau, F., Gupta, M.K., Radulescu, A., DeBosch, B., and Softic, S. (2020). Tissue-Specific Fructose Metabolism in Obesity and Diabetes. Curr Diab Rep 20, 64.

Helsley, R.N., Park, S.-H., Vekaria, H.J., Sullivan, P.G., Conroy, L.R., Sun, R.C., Romero, M.d.M., Herrero, L., Bons, J., King, C.D., et al. (2023). Ketohexokinase-C Mediates Global Protein Acetylation to Decrease Carnitine Palmitoyltransferase 1a Mediated Fatty Acid Oxidation. Journal of Hepatology. Accepted 2/9/23.

Huard, K., Ahn, K., Amor, P., Beebe, D.A., Borzilleri, K.A., Chrunyk, B.A., Coffey, S.B., Cong, Y., Conn, E.L., Culp, J.S., et al. (2017). Discovery of Fragment-Derived Small Molecules for in Vivo Inhibition of Ketohexokinase (KHK). J Med Chem 60, 7835–7849.

Inci, M.K., Park, S.H., Helsley, R.N., Attia, S.L., and Softic, S. (2022). Fructose Impairs Fat Oxidation: Implications for the Mechanism of Western diet-induced NAFLD. J Nutr Biochem, 109224.

Ishimoto, T., Lanaspa, M.A., Le, M.T., Garcia, G.E., Diggle, C.P., Maclean, P.S., Jackman, M.R., Asipu, A., Roncal-Jimenez, C.A., Kosugi, T., et al. (2012). Opposing effects of fructokinase C and A isoforms on fructose-induced metabolic syndrome in mice. Proc Natl Acad Sci U S A 109, 4320–4325.

Ishimoto, T., Lanaspa, M.A., Rivard, C.J., Roncal-Jimenez, C.A., Orlicky, D.J., Cicerchi, C., McMahan, R.H., Abdelmalek, M.F., Rosen, H.R., Jackman, M.R., et al. (2013). High-fat and high-sucrose (western) diet induces steatohepatitis that is dependent on fructokinase. Hepatology 58, 1632–1643.

Jang, C., Hui, S., Lu, W., Cowan, A.J., Morscher, R.J., Lee, G., Liu, W., Tesz, G.J., Birnbaum, M.J., and Rabinowitz, J.D. (2018). The Small Intestine Converts Dietary Fructose into Glucose and Organic Acids. Cell Metab 27, 351–361 e353.

Johnson, R.K., Appel, L.J., Brands, M., Howard, B.V., Lefevre, M., Lustig, R.H., Sacks, F., Steffen, L.M., Wylie-Rosett, J., American Heart Association Nutrition Committee of the Council on Nutrition, P.A., et al. (2009). Dietary sugars intake and cardiovascular health: a scientific statement from the American Heart Association. Circulation 120, 1011–1020.

Kawasaki, T., Akanuma, H., and Yamanouchi, T. (2002). Increased fructose concentrations in blood and urine in patients with diabetes. Diabetes Care 25, 353–357.

Kazierad, D.J., Chidsey, K., Somayaji, V.R., Bergman, A.J., Birnbaum, M.J., and Calle, R.A. (2021). Inhibition of ketohexokinase in adults with NAFLD reduces liver fat and inflammatory markers: A randomized phase 2 trial. Med (N Y) 2, 800–813 e803.

Lanaspa, M.A., Andres-Hernando, A., Orlicky, D.J., Cicerchi, C., Jang, C., Li, N., Milagres, T., Kuwabara, M., Wempe, M.F., Rabinowitz, J.D., et al. (2018). Ketohexokinase C blockade ameliorates fructose-induced metabolic dysfunction in fructose-sensitive mice. J Clin Invest 128, 2226–2238.

Lanaspa, M.A., Sanchez-Lozada, L.G., Cicerchi, C., Li, N., Roncal-Jimenez, C.A., Ishimoto, T., Le, M., Garcia, G.E., Thomas, J.B., Rivard, C.J., et al. (2012). Uric acid stimulates fructokinase and accelerates fructose metabolism in the development of fatty liver. PLoS One 7, e47948.

Lee, A.H., Scapa, E.F., Cohen, D.E., and Glimcher, L.H. (2008). Regulation of hepatic lipogenesis by the transcription factor XBP1. Science 320, 1492–1496.

Liu, L., Li, T., Liao, Y., Wang, Y., Gao, Y., Hu, H., Huang, H., Wu, F., Chen, Y.G., Xu, S., et al. (2020). Triose Kinase Controls the Lipogenic Potential of Fructose and Dietary Tolerance. Cell Metab 32, 605–618 e607.

Maryanoff, B.E., O’Neill, J.C., McComsey, D.F., Yabut, S.C., Luci, D.K., Gibbs, A.C., and Connelly, M.A. (2012). Pyrimidinopyrimidine inhibitors of ketohexokinase: exploring the ring C2 group that interacts with Asp-27B in the ligand binding pocket. Bioorg Med Chem Lett 22, 5326–5329.

Maryanoff, B.E., O’Neill, J.C., McComsey, D.F., Yabut, S.C., Luci, D.K., Jordan, A.D., Jr., Masucci, J.A., Jones, W.J., Abad, M.C., Gibbs, A.C., et al. (2011). Inhibitors of Ketohexokinase: Discovery of Pyrimidinopyrimidines with Specific Substitution that Complements the ATP-Binding Site. ACS Med Chem Lett 2, 538–543.

Moreau, F., Brunao, B.B., Liu, X.Y., Tremblay, F., Fitzgerald, K., Avila-Pacheco, J., Clish, C., Kahn, R.C., and Softic, S. (2022). Liver-specific FGFR4 knockdown in mice on a HFD increases bile acid synthesis and improves hepatic steatosis. J Lipid Res, 100324.

Nishida, C., Uauy, R., Kumanyika, S., and Shetty, P. (2004). The joint WHO/FAO expert consultation on diet, nutrition and the prevention of chronic diseases: process, product and policy implications. Public Health Nutr 7, 245–250.

Ouyang, X., Cirillo, P., Sautin, Y., McCall, S., Bruchette, J.L., Diehl, A.M., Johnson, R.J., and Abdelmalek, M.F. (2008). Fructose consumption as a risk factor for non-alcoholic fatty liver disease. J Hepatol 48, 993–999.

Park, S.H., Helsley, R.N., Fadhul, T., Willoughby, J.L.S., Noetzli, L., Tu, H.C., Solheim, M.H., Fujisaka, S., Pan, H., Dreyfuss, J.M., et al. (2023). Fructose Induced KHK-C Increases ER Stress and Modulates Hepatic Transcriptome to Drive Liver Disease in Diet-Induced and Genetic Models of NAFLD. bioRxiv.

Park, S.H., Helsley, R.N., Noetzli, L., Tu, H.C., Wallenius, K., O’Mahony, G., Boucher, J., Liu, J., and Softic, S. (2021). A luminescence-based protocol for assessing fructose metabolism via quantification of ketohexokinase enzymatic activity in mouse or human hepatocytes. STAR Protoc 2, 100731.

Phillips, M.J., Little, J.A., and Ptak, T.W. (1968). Subcellular pathology of hereditary fructose intolerance. Am J Med 44, 910–921.

Radulescu, A., Killian, M., Kang, Q., Yuan, Q., and Softic, S. (2022). Dietary Counseling Aimed at Reducing Sugar Intake Yields the Greatest Improvement in Management of Weight and Metabolic Dysfunction in Children with Obesity. Nutrients 14.

Saxena, A.R., Lyle, S.A., Khavandi, K., Qiu, R., Whitlock, M., Esler, W.P., and Kim, A.M. (2022). A phase 2a, randomized, double-blind, placebo-controlled, 3-arm, parallel-group study to assess the efficacy, safety, tolerability, and pharmacodynamics of PF-06835919 in patients with nonalcoholic fatty liver disease and type 2 diabetes mellitus. Diabetes Obes Metab.

Shepherd, E.L., Saborano, R., Northall, E., Matsuda, K., Ogino, H., Yashiro, H., Pickens, J., Feaver, R.E., Cole, B.K., Hoang, S.A., et al. (2021). Ketohexokinase inhibition improves NASH by reducing fructose-induced steatosis and fibrogenesis. JHEP Rep 3, 100217.

Singh, S.K., and Sarma, M.S. (2022). Hereditary fructose intolerance: A comprehensive review. World J Clin Pediatr 11, 321–329.

Softic, S., Boucher, J., Solheim, M.H., Fujisaka, S., Haering, M.F., Homan, E.P., Winnay, J., Perez-Atayde, A.R., and Kahn, C.R. (2016a). Lipodystrophy Due to Adipose Tissue-Specific Insulin Receptor Knockout Results in Progressive NAFLD. Diabetes 65, 2187–2200.

Softic, S., Cohen, D.E., and Kahn, C.R. (2016b). Role of Dietary Fructose and Hepatic De Novo Lipogenesis in Fatty Liver Disease. Dig Dis Sci 61, 1282–1293.

Softic, S., Gupta, M.K., Wang, G.X., Fujisaka, S., O’Neill, B.T., Rao, T.N., Willoughby, J., Harbison, C., Fitzgerald, K., Ilkayeva, O., et al. (2017). Divergent effects of glucose and fructose on hepatic lipogenesis and insulin signaling. J Clin Invest 127, 4059–4074.

Softic, S., and Kahn, C.R. (2019). Fatty liver disease: is it nonalcoholic fatty liver disease or obesity-associated fatty liver disease? Eur J Gastroenterol Hepatol 31, 143.

Softic, S., Kirby, M., Berger, N.G., Shroyer, N.F., Woods, S.C., and Kohli, R. (2012). Insulin concentration modulates hepatic lipid accumulation in mice in part via transcriptional regulation of fatty acid transport proteins. PLoS One 7, e38952.

Softic, S., Meyer, J.G., Wang, G.X., Gupta, M.K., Batista, T.M., Lauritzen, H., Fujisaka, S., Serra, D., Herrero, L., Willoughby, J., et al. (2019). Dietary Sugars Alter Hepatic Fatty Acid Oxidation via Transcriptional and Post-translational Modifications of Mitochondrial Proteins. Cell Metab 30, 735–753 e734.

Softic, S., Stanhope, K.L., Boucher, J., Divanovic, S., Lanaspa, M.A., Johnson, R.J., and Kahn, C.R. (2020). Fructose and hepatic insulin resistance. Crit Rev Clin Lab Sci, 1-15.

Sternisha, S.M., and Miller, B.G. (2019). Molecular and cellular regulation of human glucokinase. Arch Biochem Biophys 663, 199–213.

Taylor, N. (2021). Pfizer dumps midphase NASH prospect, slew of early efforts in Q2 clear-out.

Vandercammen, A., Detheux, M., and Van Schaftingen, E. (1992). Binding of sorbitol 6-phosphate and of fructose 1-phosphate to the regulatory protein of liver glucokinase. Biochem J 286 ( *Pt 1*), 253–256.

Veiga-da-Cunha, M., and Van Schaftingen, E. (2002). Identification of fructose 6-phosphate- and fructose 1-phosphate-binding residues in the regulatory protein of glucokinase. J Biol Chem 277, 8466–8473.

Young, L.E.A., Conroy, L.R., Clarke, H.A., Hawkinson, T.R., Bolton, K.E., Sanders, W.C., Chang, J.E., Webb, M.B., Alilain, W.J., Vander Kooi, C.W., et al. (2022). In situ mass spectrometry imaging reveals heterogeneous glycogen stores in human normal and cancerous tissues. EMBO Mol Med 14, e16029.

Zhao, S., Jang, C., Liu, J., Uehara, K., Gilbert, M., Izzo, L., Zeng, X., Trefely, S., Fernandez, S., Carrer, A., et al. (2020). Dietary fructose feeds hepatic lipogenesis via microbiota-derived acetate. Nature 579, 586–591.

