## supplemental data for "GalNAc-siRNA Mediated Knockdown of Ketohexokinase Versus Systemic, Small Molecule Inhibition of its Kinase Activity Exert Divergent Effects on Hepatic Metabolism in Mice on a HFD"

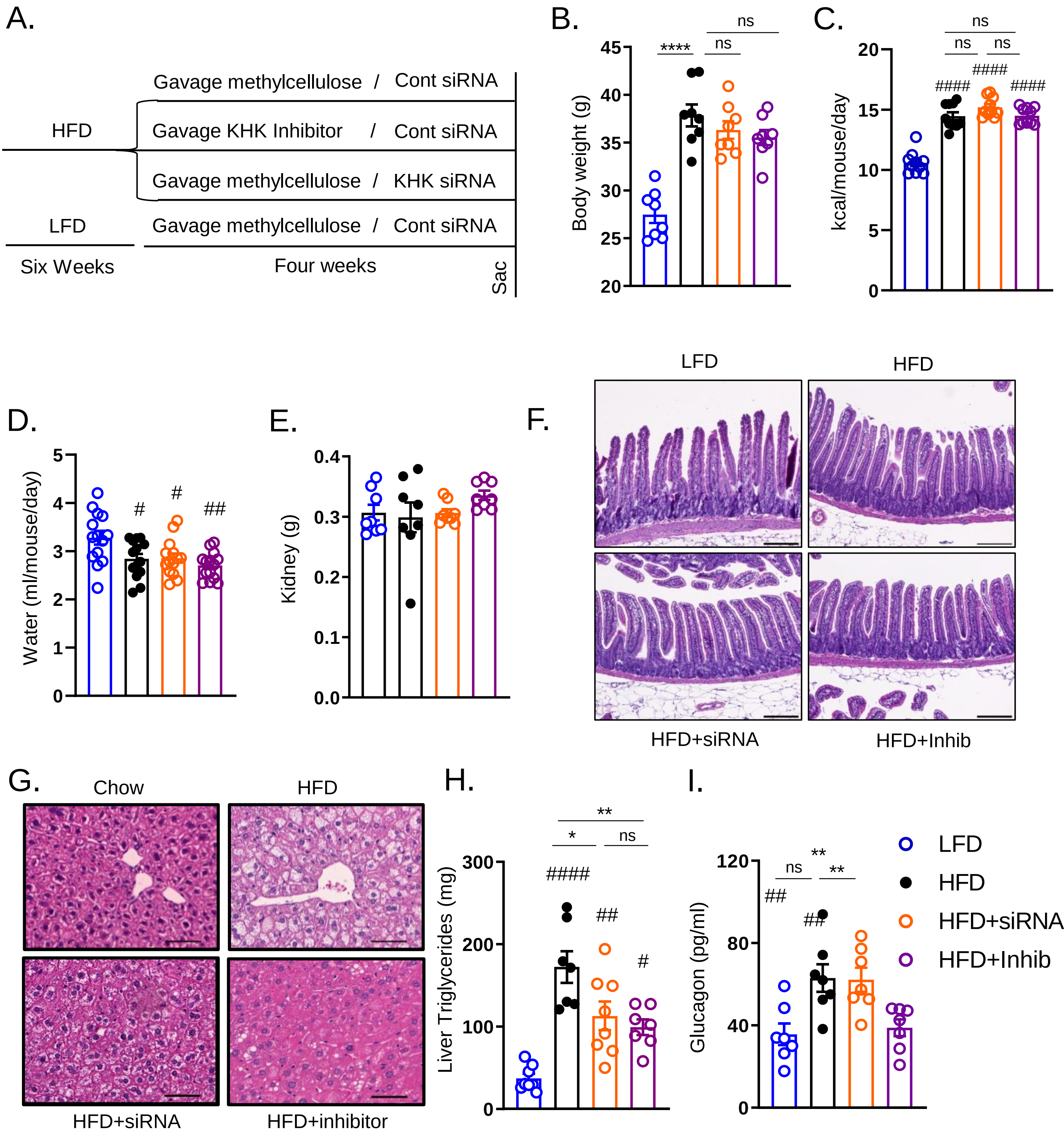


Supplemental Figure 1

A) Graphical representation of the study design. B) Body weight in these mice at the end of the study. C) Caloric intake and D) water consumption measured per cage once weekly for the duration of the experiment and averaged per mouse per day. E) Kidney weight at the time of the sacrifice. F) Representative H&E images of intestinal histology. Scale bar=200µm, G) Representative H&E images of liver histology. Scale bar=50µm H) Triglyceride accumulation in the livers of these mice. I) Serum glucagon level measured between 9-11am at sacrifice. The mice were fasted for 2h. n = 7-8 mice per group. The bar graphs show mean ±SEM. # represents statistically significant results by one way ANOVA compared to the LFD group. * represents post-hock t-test between the groups.

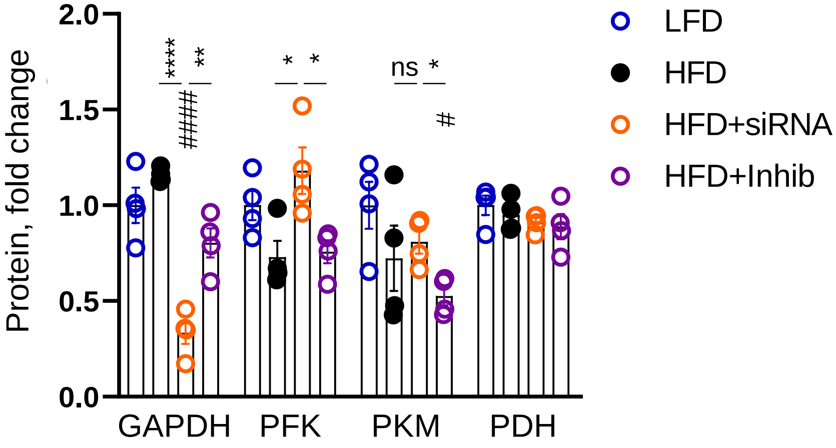


Supplemental Figure 2
Densitometry quantification of western blot data in figure 2G using Image J. n = four mice per group. The bar graphs show mean ±SEM. # represents statistically significant results by one way ANOVA compared to the LFD group. * represents post-hock t-test between the groups.

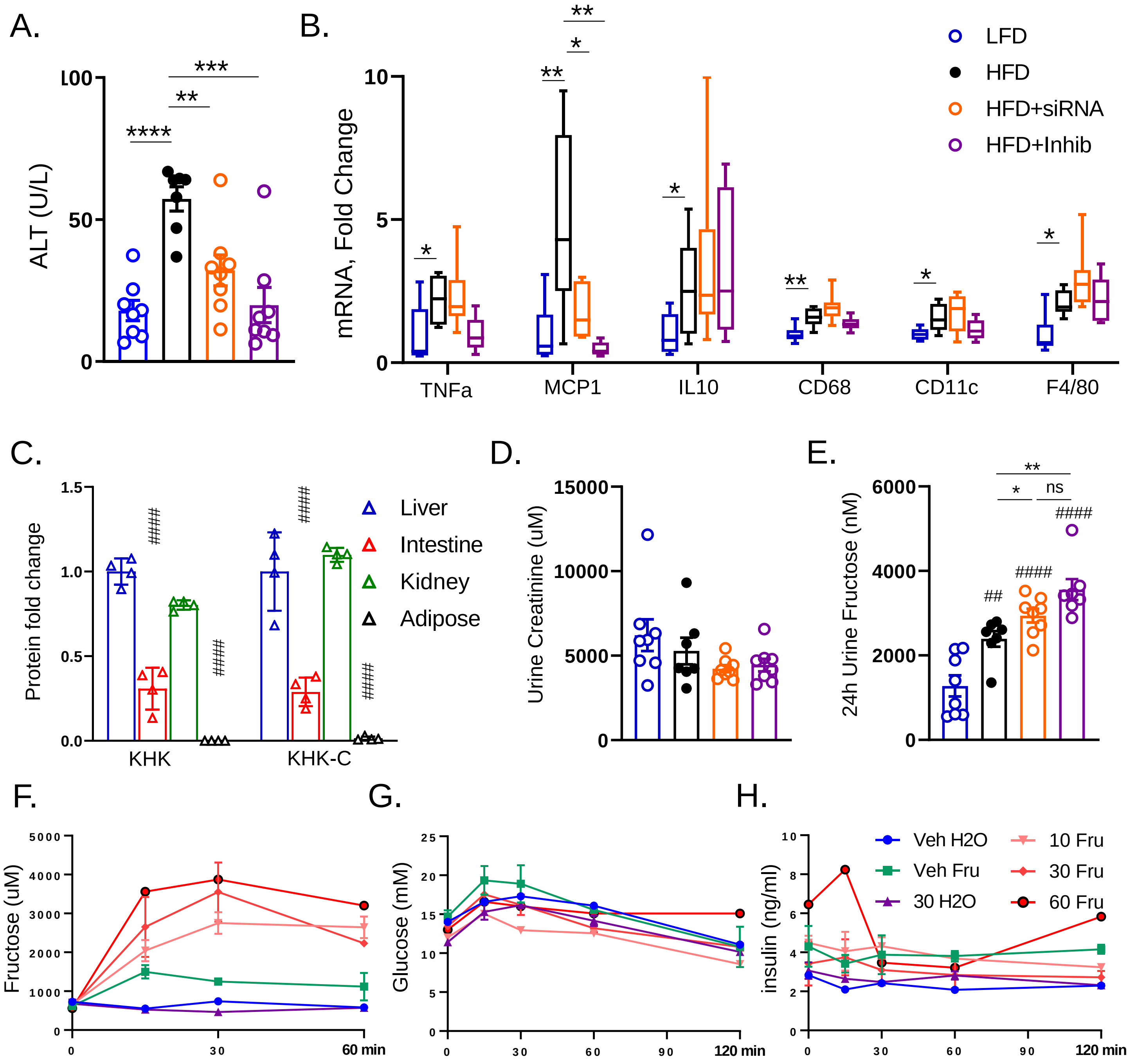


Supplemental Figure 3

A) Serum ALT at the time of the sacrifice. B) Hepatic mRNA expression of inflammatory genes in these mice. n = 6 mice per group. C) Densitometry quantification of western blot data in figure 3I using Image J. n = four mice per group. D) Urine creatinine concentration and E) Fructose concentration in the urine collected over 24h between 9-10 weeks on the diet. n= 7-8 mice per group. The bar graphs show mean ±SEM. # represents statistically significant results by one way ANOVA compared to the LFD group. * represents post-hock t-test between the groups. Serum F) fructose, G) glucose and G) insulin levels in C57Bl/6J male mice gavaged with 10 mg/kg, 30 mg/kg or 60 mg/kg of the ketohexokinase inhibitor or methylcellulose vehicle one hour before 6 mg/kg fructose or water gavage. n = 1-2 mice per group. The data points represent mean ±SEM.

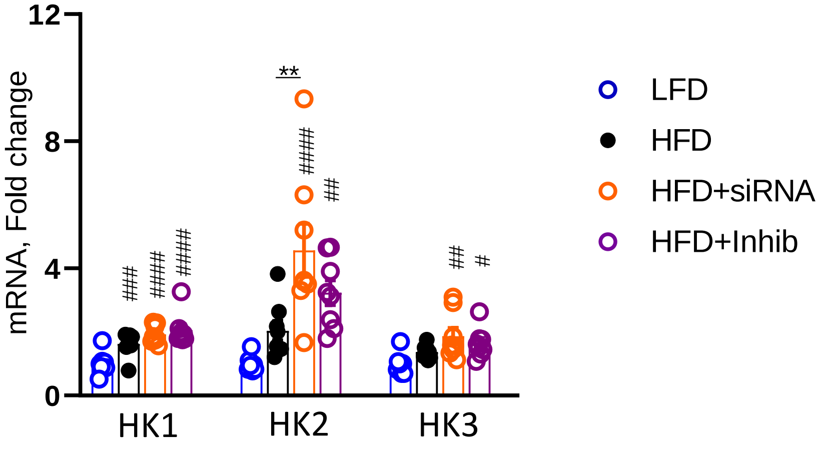


Supplemental Figure 4
Hepatic mRNA expression of hexokinase isoforms from the livers of mice obtained at the sacrifice. n = 6 mice per group. The bar graphs show mean ±SEM. # represents statistically significant results by one way ANOVA compared to the LFD group. * represents post-hock t-test between the groups.

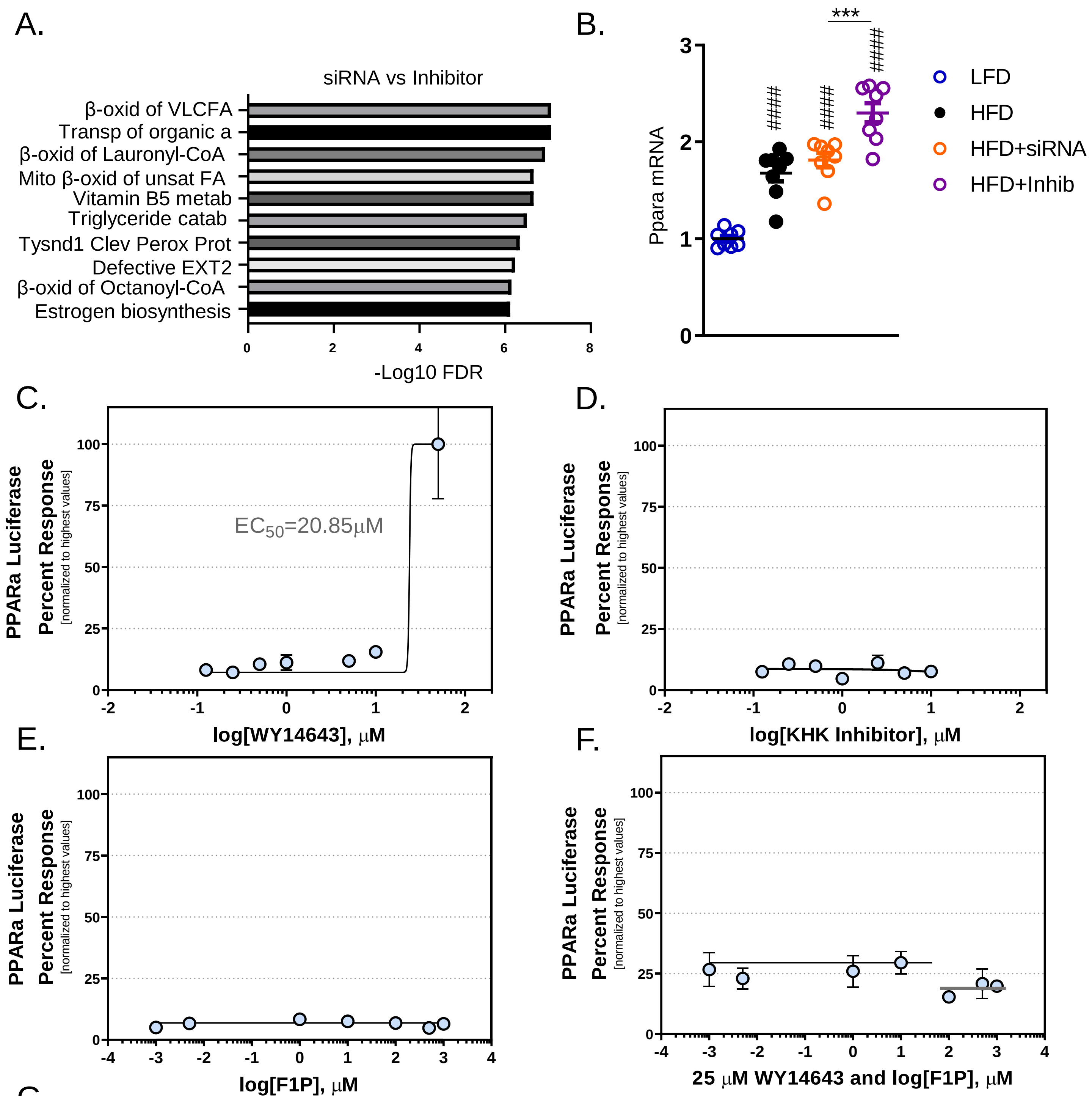


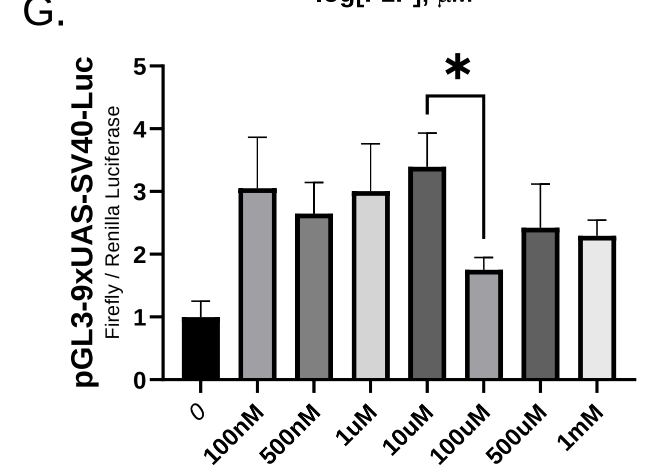


Supplemental Figure 5
A) Reactome pathway analysis using RNAseq data. B) Hepatic mRNA expression of PPARa. n = 6 mice per group. PPARa luciferase reporter assay following treatment with increasing concentrations of C) WY14643, D) KHK inhibitor and E) Fructose-1 phosphate (F1P). F) PPARa luciferase reporter assay following treatment with 25 uM of WY14643 and increasing concentrations of F1P. n = 6 wells per condition. G) bargraph representation of the data in figure F. The bar graphs show mean ±SEM. *statistically significant result by one way ANOVA.

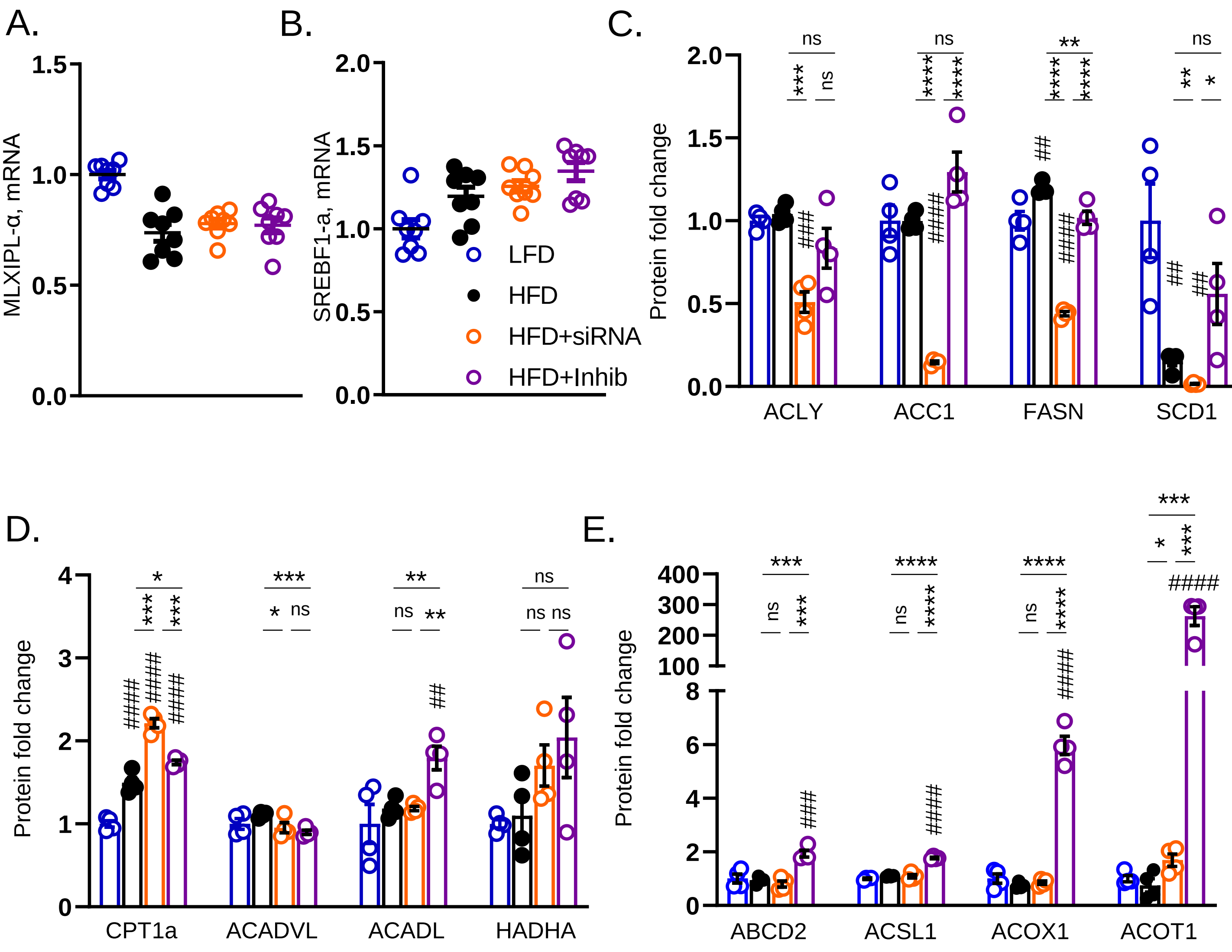


Supplemental Figure 6
Hepatic mRNA expression of A) Chrebp and B) Srebp1c. n = 6 mice per group. Densitometry quantification of western blot data using Image J C) from figure 3I, D) figure 6F and E) figure 6H. n = four mice per group. The bar graphs show mean ±SEM. # represents statistically significant results by one way ANOVA compared to the LFD group. * represents post-hock t-test between the groups.

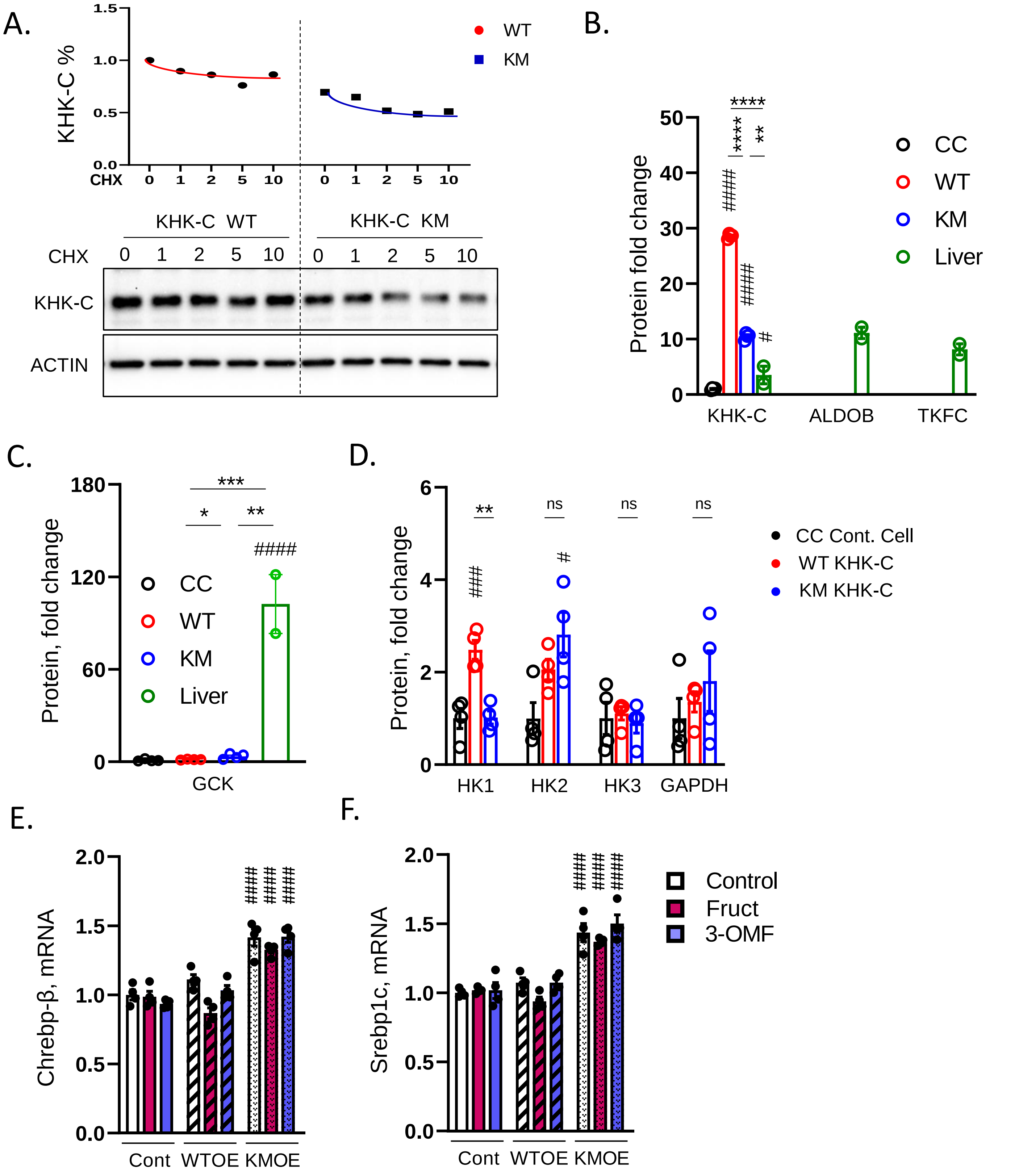


Supplemental Figure 7

A) Western blot of KHK-C from wild type (WT) or kinase dead mutant (KM) KHK-C overexpressing cells treated with increasing doses of cycloheximide (CHX). The graph above represents Image J quantification of the data. Densitometry quantification of western blot data using Image J from B) figure 7B, C) figure 7K and D) figure 7K. n = four mice per group. mRNA expression of Chrebf and F) Srebp1c. n=4 wells per group. The bar graphs show mean ±SEM. # represents statistically significant results by one way ANOVA compared to the control group. * represents post-hock t-test.
